# STAT3 signaling in B cells controls germinal center zone organization and recycling

**DOI:** 10.1101/2022.08.12.503811

**Authors:** Adam J Fike, Sathi Babu Chodisetti, Nathaniel E Wright, Kristen N Bricker, Phillip P Domeier, Mark Maienschein-Cline, Aaron M Rosenfeld, Sara A Luckenbill, Julia L Weber, Nicholas M Choi, Eline T Luning Prak, Malay Mandal, Marcus R Clark, Ziaur SM Rahman

## Abstract

Germinal centers (GCs), sites of antibody affinity maturation, are organized into dark (DZ) and light (LZ) zones. Here, we uncovered a B cell intrinsic role for STAT3 in GC DZ and LZ organization. Altered zonal organization of STAT3-deficient GCs dampened GC output of long-lived plasma cells (LL-PCs) but increased memory B cells (MBCs). Tfh-GC B cell interaction drive STAT3 tyrosine 705 and serine 727 phosphorylation in LZ B cells, facilitating their recycling into the DZ. An inducible system confirmed STAT3 is not involved in initiating or maintaining the GC but sustains GC zonal organization by regulating GC B cell recycling. RNAseq and ChIPseq analysis identified genes regulated by STAT3 that are critical for LZ cell recycling and transiting through the DZ proliferation and differentiation phases of the DZ. Thus, STAT3 signaling in B cells controls GC zone organization and recycling, and GC egress of LL-PCs, but negatively regulates MBC output.

**Summary:** Fike et al. describe a previously unrecognized mechanism by which B cell intrinsic STAT3 signaling controls the germinal center (GC) dark and light zone organization, GC B cell recycling, and GC output of long-lived plasma cells but negatively regulates memory B cells.

## INTRODUCTION

Germinal centers (GCs) are specialized microenvironments that generate high-affinity antibodies for immune protection (De Silva and Klein, 2015; Laidlaw and Ellebedy, 2022; Victora and Nussenzweig, 2012; Young and Brink, 2021). GCs are canonically organized into two distinct niches, the dark (DZ) and light (LZ) zone (Dominguez-Sola et al., 2015; Sander et al., 2015; Victora and Nussenzweig, 2012; Young and Brink, 2021) which we define as ‘GC zone organization’. The DZ can be subdivided into the DZ proliferation (DZp) and the DZ differentiation (DZd) where GC B cells undergo somatic hypermutation (SHM) of the immunoglobulin (Ig) genes (Kennedy and Clark, 2021; Kennedy et al., 2020). In the LZ, rich in follicular dendritic cells (FDCs) and follicular helper (Tfh) T cells, GC B cells engage antigen via their B cell receptor (BCR) from FDC-captured or soluble antigens and undergo Tfh-driven selection by presenting peptides via MHC-II (Mesin et al., 2016; Shlomchik et al., 2019; Young and Brink, 2021). After selection, LZ B cells either recycle back to the DZ for further proliferation and SHM of Ig genes, differentiate into long-lived plasma (LL-PCs) or memory B cells (MBCs), or undergo apoptosis. Tight regulation of GC DZ and LZ is important for the generation of high-affinity antibody-producing LL-PCs and MBCs. The mechanisms controlling GC zonal compartmentalization and GC output are incompletely defined.

Current models suggest that the magnitude of B cell receptor and T cell signals dictate the B cell fate to either exit the GC or remain in the GC and initiate cyclic reentry (Inoue et al., 2021; Ise et al., 2018; Laidlaw and Cyster, 2021). B cells with weak BCR and poor T cell help undergo apoptosis. Weak T cell help (e.g., limited CD40, IL-4, and/or IL-21), but sufficient BCR stimulation leads to MAPK-mediated repression of BCL-6 and GC exit as MBCs through the cooperation of several transcription factors, including BACH2, HHEX, TLE3, and SKI (Laidlaw et al., 2020; Shinnakasu and Kurosaki, 2017). An intermediate level of T cell help induces Myc and FOXO1 expression/activation resulting in LZ cell recycling into the DZ (Dominguez-Sola et al., 2015; Dominguez-Sola et al., 2012; Inoue et al., 2017; Roberto et al., 2021; Sander et al., 2015). Myc transcriptionally regulates AP4 and UHRF1 in concert with MIZ1 to allow cell cycle progression (Chou et al., 2016; Toboso-Navasa et al., 2020). FOXO1 expression is mediated by mTORC1, TGFβ, and/or IL-10 and its nuclear translocation is dependent on reduced PI3K activity through pAkt (Albright et al., 2019; Dominguez-Sola et al., 2015; Ersching et al., 2017; Inoue et al., 2017; Laidlaw et al., 2017; Sander et al., 2015). FOXO1 regulates expression of *Cxcr4*, encoding a chemokine receptor responsive to the CXCL12 gradient to promote LZ cell migration to the DZ (Dominguez-Sola et al., 2015). Several other signals support various aspects of DZ, including proliferation, cell cycle, and metabolism (Ersching et al., 2017; Inoue et al., 2021; Pae et al., 2021). Finally, strong T cell help results in IRF4-mediated repression of BCL-6, induction of BLIMP1(*Prdm1*), and GC exit as plasma cells (Shinnakasu and Kurosaki, 2017; Xu et al., 2015; Zhang et al., 2018). These data suggest that GC B cell:Tfh interactions in the LZ orchestrate a complex interplay of signals directing the trajectory of B cell differentiation.

Signal transducer and activator of transcription 3 (STAT3) mediates signaling downstream of several stimuli important for GC biology including, but not limited to IL-6, IL-10, IL-21, Type I interferon, and mTORC1 (Arkatkar et al., 2017; Ersching et al., 2017; Hanissian and Geha, 1997; Laidlaw et al., 2017; Ray et al., 2014; Tangye and Ma, 2020; Xu et al., 2017). Maximal STAT3 activity is regulated through the phosphorylation of tyrosine 705 and serine 727, followed by dimerization and nuclear translocation for target gene transcription (Wen et al., 1995). Several studies have described the roles of upstream cytokine receptors, especially the IL-21 receptor (IL-21R), in GC and high-affinity antibody responses (Avery et al., 2010; Linterman et al., 2010; Zotos et al., 2010). While GC prematurely collapsed in the absence of IL-21R signaling in B cells (Linterman et al., 2010; Zotos et al., 2010), STAT3 deficiency in B cells had no significant effect on mounting a peak GC response albeit attenuated plasma cell response (ref). These data indicate that the mechanisms by which STAT3, which can be downstream of various signals, regulate GC and long-lived antibody responses do not entirely recapitulate IL-21 mediated regulation of GC and plasma cell responses.

Here we used several developmental stage-specific B cell conditional deletion systems to discover a previously unrecognized role for STAT3 signaling in B cells in the organization of the GC DZ and LZ. We identified that STAT3 in B cells is required for GC B cell differentiation into LL-PCs but functions as a negative regulator of MBC formation. However, altered DZ and LZ organization in the absence of STAT3 in B cells exhibited minimal effects on SHM of Ig genes and clonal diversity of the NP-specific GC response. Tfh cell-derived signals phosphorylated STAT3 Tyrosine 705 (pY705) and Serine 727 (pS727) residues in LZ B cells to drive GC cell recycling. Using temporally controlled deletion system, we found that B cell intrinsic STAT3 is not required for GC initiation or maintenance but is important for sustaining GC DZ and LZ compartmentalization. The alteration in the zonal distribution of STAT3-deficient GC B cells was not the result of substantial defects in GC B cell proliferation or apoptosis.

Finally, we identified the transcriptional program elicited by STAT3 in GC B cells, including genes critically involved in GC B cell transition from the LZ through the DZ proliferation (DZp) and differentiation (DZd) phases. Together, these data highlight a B cell-intrinsic role of STAT3 in positive regulation of GC cell recycling, organization of the GC zones and the output of LL-PCs, whereas in negative regulation of MBC formation.

## RESULTS

### STAT3 signaling in B cells regulates GC dark and light zone organization

To delineate the mechanisms by which STAT3 signaling in B cells regulates the GC response, we crossed STAT3^fl/fl^ mice with CD19^Cre^ mice to conditionally delete STAT3 during the pro-B cell stage of B cell development. To mount a robust GC reaction, we employed a prime-boost strategy of immunizing mice with NP-KLH in complete freund’s adjuvant (CFA) and incomplete freund’s adjuvant (IFA) on d0 and d7, respectively. We observed no differences in total GC or Tfh responses between STAT3^fl/fl^ and STAT3^fl/fl^CD19^Cre^ mice on 14d post-NP-KLH immunization (Fig. 1A, B). However, analysis of the zonal distribution of GC B cells revealed a reduction in the frequency of DZ B cells (Fig. 1C, D) and an increased frequency of LZ B cells (Fig. 1C, E) in STAT3^fl/fl^CD19^Cre^ mice compared to control mice. The altered ratio of the GC DZ and LZ in STAT3^fl/fl^CD19^Cre^ mice strongly correlated with reduced high (NP_4_) and total (NP_29_) NP-specific IgG1 Ab responses (Fig. 1F) in the serum. We then crossed STAT3^fl/fl^ mice with CD23^Cre^ mice to delete STAT3 in peripheral B cells (Chodisetti et al., 2020b; Kwon et al., 2008). STAT3^fl/fl^CD23^Cre^ mice, immunized as described above, also had similar GC and Tfh responses to CD23^Cre^ control mice (Fig. 1G-I), but alterations to GC zone organization (reduced DZ and increased LZ B cell responses) were present (Fig. 1J-M). Staining of FDC (FDC-M1), DZ (GL-7), and LZ B cells (CD86) confirmed alterations to the LZ/DZ distribution in GCs of STAT3^fl/fl^CD23^Cre^ mice (Fig. 1N). Total CD138^+^TACI^+^ plasma cell (PC) responses were not different between the two strains (Fig. 1O); however, STAT3^fl/fl^CD23^Cre^ mice had reduced percentages and numbers of NP-specific subpopulations of PCs, including antibody-forming cells (AFCs) (Fig. 1P-S). The reduction in NP-specific AFC responses resulted in dampened serum NP-specific IgG and IgG1 Ab responses (Fig. 1T, U).

**Fig. 1.**
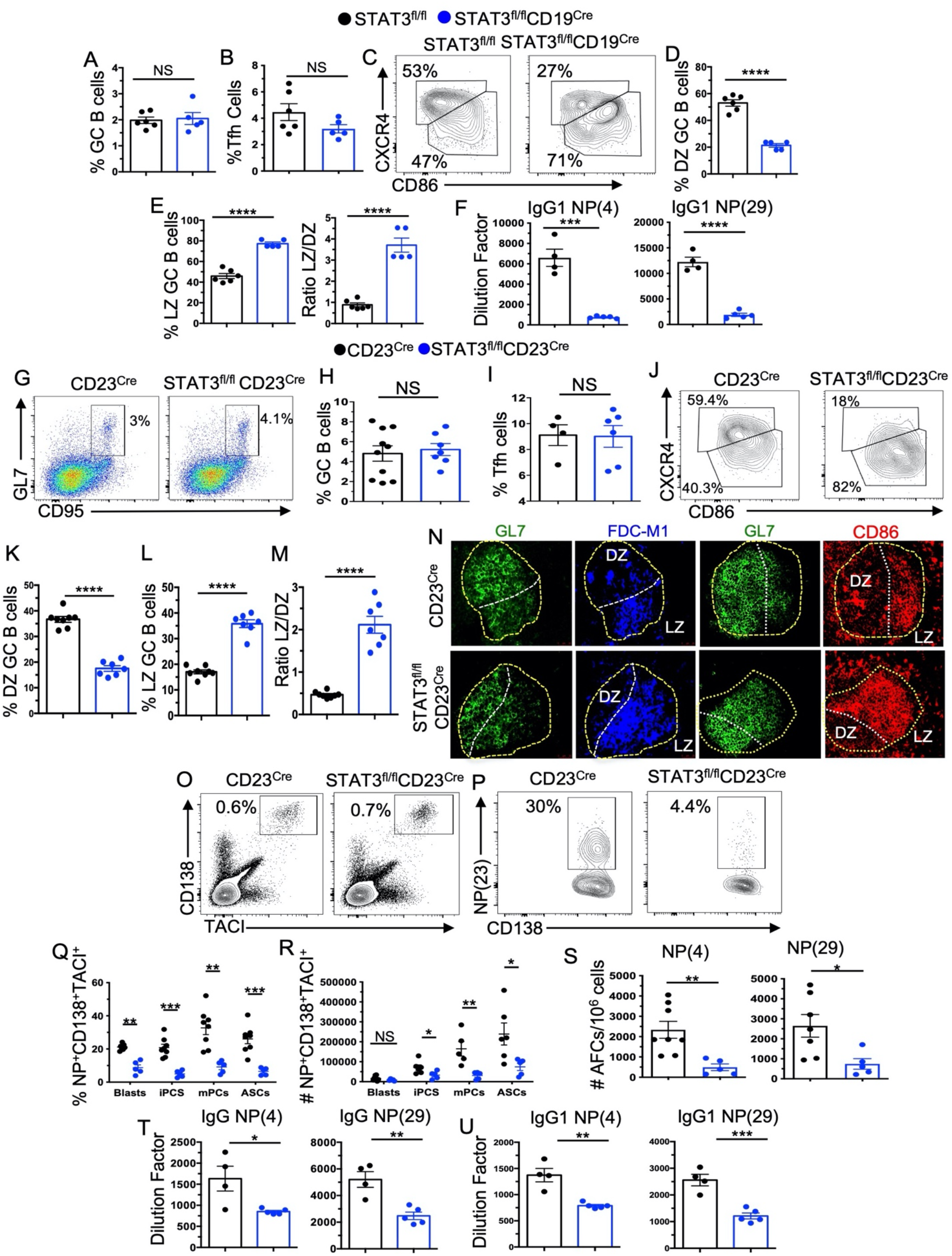
STAT3 signaling in B cells is required for GC DZ and LZ organization. **(A-F)**STAT3^fl/fl^CD19^Cre^ and STAT3^fl/fl^ control mice were immunized with NP-KLH as described in Materials and Methods. The splenic GC response was analyzed on 14d post-immunization. **(A)** Frequency of B220^+^GL-7^+^CD95^+^ GC B cells. **(B)** Frequency of CD4^+^CD44^+^CD62L^-^CXCR5^+^PD-1^+^ splenic Tfh. **(C-F)** Representative flow plots and quantification of CXCR4^hi^CD86^lo^ DZ and CXCR4^lo^CD86^hi^ LZ GC B cells. **(G)** NP(4) and NP(29) specific serum IgG1titers. **(G-M)** Flow cytometric analysis of GC responses on 14d post-NP-KLH immunized CD23^Cre^ and STAT3^fl/fl^CD23^Cre^ mice. **(G)** Gating strategy for B220^+^GL7^hi^CD95^hi^ GC B cells. Percentages of GC B cells in total B220^+^ B cells **(H)** and CD4^+^CD44^+^CD62L^-^PD-1^hi^CXCR5^hi^ Tfh **(I). (J)** Representative FACS plots for CXCR4^hi^CD86^lo^ DZ and CXCR4^lo^CD86^hi^ LZ GC B cells from indicated mice. Percentages of DZ **(K)** and LZ **(L)** GC B cells and the ratio of LZ/DZ B cells **(M)**. (**N**) Splenic sections from immunized mice were stained with GL7 (green) and FDC-M1 (blue) or anti-CD86 Ab to define (shown in dotted lines) DZ and LZ areas within the GC (200X). **(O)** Representative flow plot of total IgD^-^CD138^+^TACI^+^ plasma cells. **(P)** Representative flow plot of NP^+^IgD^-^CD138^+^TACI^+^ plasma cells. (**Q, R**) Percentages and numbers of subsets of NP-specific plasma cells (IgD^-^NP^+^CD138^+^TACI^+^B220^int^CD19^+^ blasts, IgD^-^NP^+^CD138^+^TACI^+^B220^-^CD19^+^ immature plasma cells-iPCs, IgD^-^NP^+^CD138^+^TACI^+^B220^-^CD19^-^ mature/resting plasma cells-mPCs and antibody-secreting cells-ASCs). (**S**) Numbers of NP4- and NP29-specific splenic AFCs enumerated by ELISpot. (**T, U**) NP4- and NP29-specific serum IgG and IgG1 titers measured by ELISA. Each symbol represents an individual mouse (n=5-10 mice per group) and data are presented as means ± SEM. Data in each panel represent two-four experiments. P values were calculated via an unpaired t-test or Mann-Whitney test (NS, not significant, p>0.05, *, p <0.05, **, p <0.01, ***, p <0.001, ****, p <0.0001).

To examine whether B cell-intrinsic STAT3 deficiency altered the GC zone organization early during its formation, we evaluated responses at d7 post-NP-KLH/CFA immunization without boosting. Similar to d14, the overall GC, CD4^+^ effector T cell, Tfh, and PC responses were not different from control mice (Fig. S1A, F, G, H). However, STAT3^fl/fl^CD23^Cre^ mice had altered DZ and LZ B cell frequencies and ratios (Fig. S1B-E) and reduced NP-specific AFC responses at this early stage of the GC response (Fig. S1I).

To determine whether T cell intrinsic STAT3 impacted GC architecture, we deleted STAT3 in T cells (STAT3^fl/fl^CD4^Cre^). NP-KLH-immunized STAT3^fl/fl^CD4^Cre^ mice had reduced overall GC and Tfh responses (Fig. S2A, E, F), but GC DZ/LZ organization was normal (Fig. S2B-D). Together, these data suggest that although STAT3 signaling in B cells is dispensable for mounting a quantitatively normal GC response, qualitatively it controls the GC DZ and LZ organization which is important for PC differentiation and IgG Ab production.

### STAT3 deletion in GC B cells recapitulates alteration in the GC DZ and LZ organization

We found that STAT3 expression is upregulated in GC B cells relative to non-GC B cells (Fig. 2A). Therefore, we crossed STAT3^fl/fl^ mice with Cγ1^Cre^ mice such that STAT3 would be deleted in IgG1 class-switched B cells during the initiation of the GC (Casola et al., 2006).

**Fig. 2.**
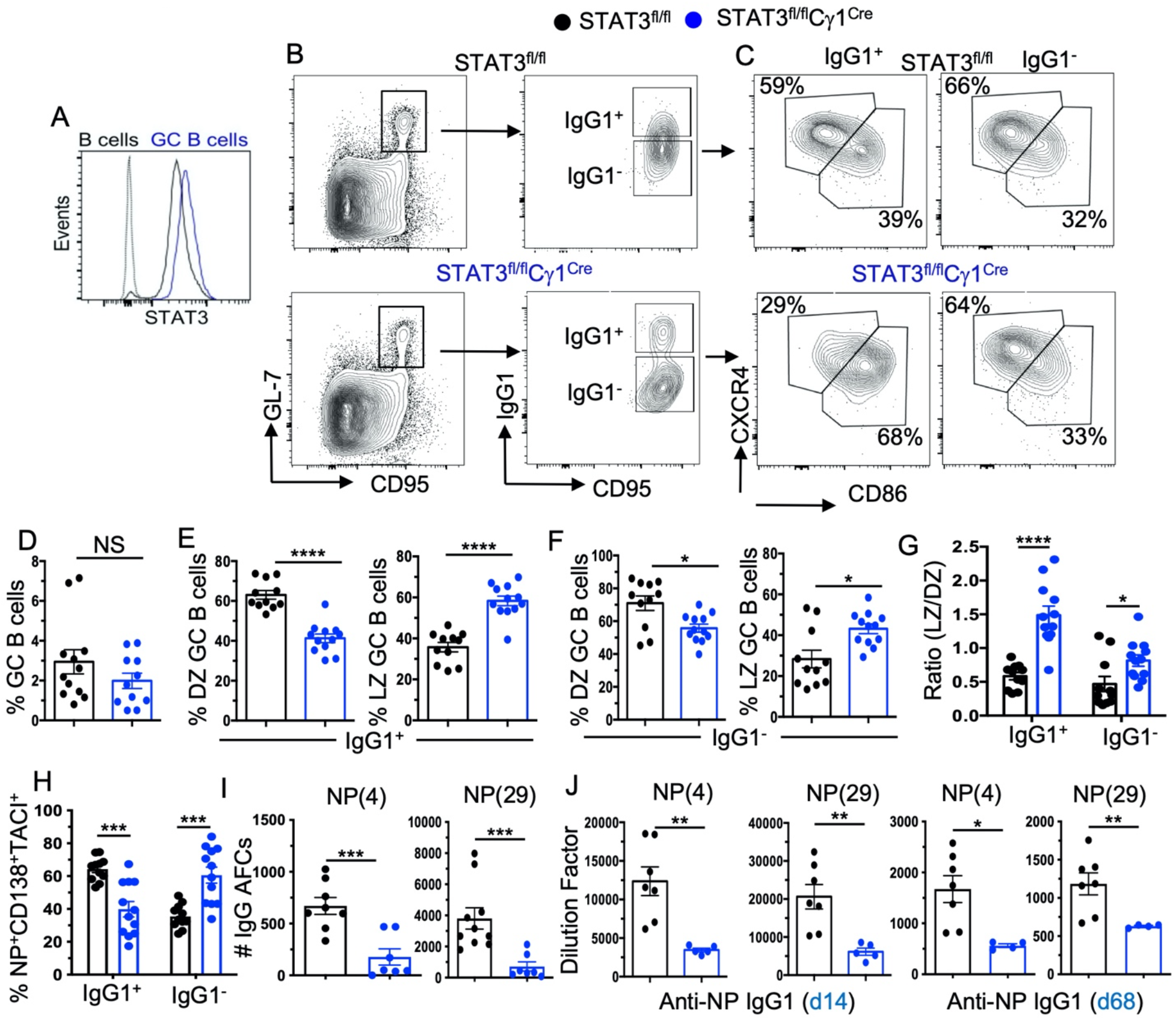
STAT3 deletion in post-CSR B cells recapitulates GC DZ and LZ disorganization. (**A**) Representative histogram of STAT3 expression in B220^+^GL-7^+^CD95^+^ GC B cells (blue) compared to B220^+^GL-7^-^CD95^-^ non-GC B cells (black) and an isotype control (gray). (**B, C**) Gating strategy of IgG1^+^ and IgG1^-^ GC B cells and their respective LZ and DZ cells in STAT3^fl/fl^C±1^Cre^ and STAT3^fl/fl^ control mice 14d post-immunization with NP-KLH. (**D**) Flow cytometric quantification of the GC response in indicated mice. (**E-G**) Quantification of DZ and LZ populations from IgG1^+^ or IgG1^-^ GC B cells. (**H**) Percentages of IgG1^+^ and IgG1^-^ IgD^-^ NP^+^CD138^+^TACI^+^ plasma cells from indicated mice. (**I**) Numbers of NP4- and NP29-specific splenic AFCs enumerated by ELISpot. (**J**) NP4- and NP-29 specific IgG1 serum titers measured by ELISA on 14d and 68d post-immunization. Each symbol represents an individual mouse and data are presented as means ±SEM (n=5-12 mice per group). Data in each panel represent two-three experiments. P values were calculated via an unpaired t-test or Mann-Whitney test (NS, p>0.05, *, p <0.05, **, p <0.01, ***, p <0.001, ****, p <0.0001).

Although the overall GC (Fig. 2B) and Tfh (not shown) responses in STAT3^fl/fl^Cγ1^Cre^ and STAT3^fl/fl^ control mice were similar, IgG1^+^ GC B cells in STAT3^fl/fl^Cγ1^Cre^ mice had significantly reduced and increased percentages of DZ and LZ B cells, respectively (Fig. 2C-G). The overall CD138^+^TACI^+^ splenic PC responses in STAT3^fl/fl^Cγ1^Cre^ and STAT3^fl/fl^ mice did not differ (Fig. 2H); however, numbers of NP4- and NP29-specific IgG1 AFCs in STAT3^fl/fl^Cγ1^Cre^ mice were lower than STAT3^fl/fl^ control mice (Fig. 2I). The GC DZ/LZ disorganization and reduced NP-specific AFC responses in STAT3^fl/fl^Cγ1^Cre^ mice were associated with dampened serum antibody titers at peak (d14) and late (d68) timepoints (Fig. 2J). These data demonstrate a GC B cell-specific role for STAT3 in the regulation of GC DZ and LZ organization and, in part, GC output of PCs and IgG antibodies.

### STAT3 is important for the GC zone organization during influenza viral infection

We next infected STAT3^fl/fl^CD23^Cre^ and CD23^Cre^ control mice intranasally with influenza virus to determine whether STAT3 also regulates GC DZ and LZ organization during a viral infection. We found an increase in overall GC responses in the mediastinal draining lymph nodes (mLN) of STAT3^fl/fl^CD23^Cre^ compared to CD23^Cre^ mice 14d post-infection (Fig. 3A). No differences in weight loss were observed between STAT3^fl/fl^CD23^Cre^ and CD23^Cre^ infected mice (data not shown). In addition, we observed reduced DZ and increased LZ B cell responses in STAT3^fl/fl^CD23^Cre^ mice compared to CD23^Cre^ control mice (Fig. 3B-E). STAT3^fl/fl^CD23^Cre^ mice had reduced CD138^+^TACI^+^ PC and virus-specific IgG Ab responses than CD23^Cre^ control mice (Fig. 3F-G). These findings were recapitulated in influenza virus infected STAT3^fl/fl^Cγ1^Cre^ and control mice. The ratio of LZ to DZ B cells was increased in IgG1^+^ GC B cells (Fig. 3I-L), despite there being no differences in overall GC responses between the two strains (Fig. 3H). STAT3^fl/fl^Cγ1^Cre^ mice also had reduced CD138^+^TACI^+^ PC and virus-specific IgG Ab responses than control mice (Fig. 3M, N). Together, data from two types of immunization (prime-boost vs single) and infection models utilizing total (CD23^Cre^ or CD19^Cre^) and early GC (Cγ1^Cre^) B cell-specific conditional STAT3 KO mouse systems indicate that STAT3 signaling in B cells is required for GC DZ and LZ organization and prolonged class-switched IgG antibodies.

**Fig. 3.**
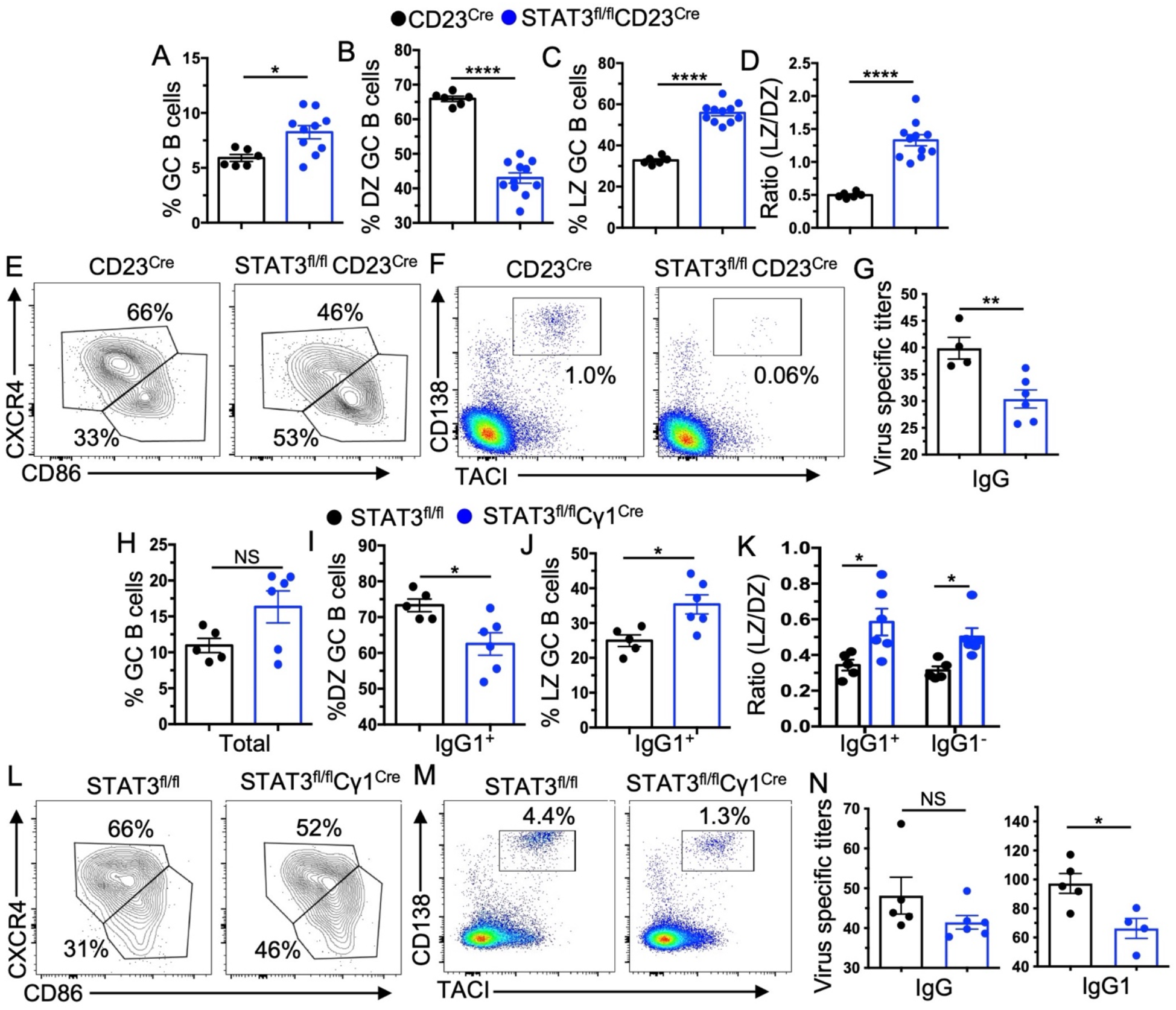
B cell intrinsic STAT3 is important for the GC organization during viral infection. (**A-G**) Assessment of B cell responses in the draining mediastinal lymph node (mLN) on 14d post-influenza viral infection of CD23^Cre^ and STAT3^fl/fl^CD23^Cre^ mice. **(A)** Percentage of B220^+^GL-7^+^CD95^+^ GC B cells. (**B-D**) Quantification of DZ, LZ, and LZ/DZ ratio of GC B cells. (**E**) Representative flow plots depicting LZ and DZ GC B cells. (**F**) Representative flow plots of IgD^-^CD138^+^TACI^+^ plasma cells in mLNs. (**G**) Anti-influenza virus specific serum IgG titers measured by ELISA. (**H-N**) STAT3^fl/fl^ and STAT3^fl/fl^C1^cre^ mice were infected with influenza virus and B cell responses were interrogated on 14d post-infection in mLNs. (**H-K**) Percentage of total GC B cells and IgG1^+^ and IgG1^-^ LZ/DZ cells. (**L**) Representative flow plots of LZ and DZ cells from IgG1^+^ GC B cells. (**M**) Representative flow plots of IgD^-^CD138^+^TACI^+^IgG1^+^ plasma cells. (**N**) Quantification of serum influenza virus-specific IgG and IgG1 titers. Each symbol represents an individual mouse and data are presented as means ±SEM (n=5-10 mice per group). Accumulative data represent two-three experiments. P values were calculated via an unpaired t-test or Mann-Whitney test (NS, p>0.05, *, p <0.05, **, p <0.01, ***, p <0.001, ****, p <0.0001).

### DZ defect caused by STAT3 deficiency in B cells has modest effects on SHM and the antibody repertoire

To determine whether GC zone disorganization mediated by STAT3 deficiency in B cells affects SHM, we amplified and sequenced the NP-specific VH186.2 segment from FACS-sorted NP^+^ GC B cells from STAT3^fl/fl^CD23^Cre^ and CD23^Cre^ control mice 14d post-immunization. In line with previous reports (Dominguez-Sola et al., 2015; Sander et al., 2015), despite defect in GC zone distribution we found no effect of STAT3 deletion on the number or location of mutations within VH186.2 (Fig. S3A). No difference in the frequency of W33L mutation as evidence for affinity maturation was observed (Allen et al., 1988) (Fig. S3B).

V-gene mutations accumulate throughout the persistence of the GC reaction (Kuraoka et al., 2016). Given the lack of difference in SHM in VH186.2 on d14 as assessed through low-throughput cloning, we sought to comprehensively evaluate the effects of STAT3 deficiency by sequencing the VH genes from FACS-sorted NP-specific and non-NP-specific GC B cells from STAT3^fl/fl^CD23^Cre^ and CD23^Cre^ control mice 21d post-NP-KLH immunization. We identified similar clone counts, CDR3 lengths, and Igh V gene identity in STAT3 sufficient and deficient animals including when stratified by their NP binding status (Fig. S3C). To determine if STAT3 deficiency influenced the degree of clonal expansion, we analyzed the distribution of copy number fractions (Fig. S3D) which showed slightly more total reads (high copy), but fewer clones among the STAT3 sufficient NP-binders. Analysis of Vh gene usage revealed significant skewing towards Vh1-72, consistent with NP-binding B cells (Fig. S3E) (Kuraoka et al., 2016), but Vh gene usage did not differ between STAT3 sufficient and deficient GC B cells. To determine if STAT3 deficiency altered the level of somatic mutation (SHM), we compared the fraction of clones with SHM levels of 2% or greater (based on the fraction of nucleotides that differ in each clone’s Vh gene, compared to its nearest germline gene, counting each clone only once). The fraction of mutated clones was similar in all four groups, but the level of SHM among mutated clones was lower in the NP-binders than in the NP-non-binders (Fig. S3E, F). Together, our findings demonstrate that the reduction in the DZ and an increase in the LZ caused by STAT3 deficiency in B cells is associated with modest alterations to the antibody repertoire.

### Defects in proliferation or cell cycle progression may not be the major driver of reduced DZ in STAT3 deficient GCs

Based on altered zonal compartmentalization of GCs in the absence of STAT3, we investigated whether reduced proliferation or increased cell death accounted for skewed DZ and LZ ratios. Percentages of Ki67^+^ cells in the DZ and LZ were similar between STAT3^fl/fl^CD23^Cre^ and CD23^Cre^ control mice 14d post-immunization (Fig. S4A). *In vivo* BrdU incorporation also revealed similar levels of proliferating GC B cells (Fig. S4B-E). Using a BrdU/EdU double pulse, we assessed GC B cell progression through S phase of the cell cycle at 14d post-immunization (Gitlin et al., 2014). The percentage of post-S phase GC B cells was lower in STAT3^fl/fl^CD23^Cre^ mice than CD23^Cre^ control mice, although no difference in the percentages of GC B cells in the early and mid S phases of the cell cycle (Fig. S4F-I). Cell cycle progression was confirmed by quantifying DNA content via Hoechst staining (Fig. S4J). All phases of the cell cycle were equivalent between CD23^Cre^ and STAT3^fl/fl^CD23^Cre^ mice except we found a reduction in the percentage of LZ GC B cells that were in S phase in the STAT3^fl/fl^CD23^Cre^ GC B cells (Fig. S4K). We found no difference in the percentages of DZ and LZ GC B cells undergoing apoptosis or cell death between the two strains (Fig. S4K). As a proxy for DNA damage, we measured the expression of serine 139 phosphorylated H2AX (S139H2AX) in DZ and LZ GC B cells and found no significant differences (Fig. S4L). We also measured apoptosis by histological SR-FLICA staining, which detects active caspases 3 and 7, involved in apoptosis induction, and found no difference (Fig. S4M, N). We repeated the BrdU/EdU double pulse experiment during the early stage of the GC response (d7) following a single immunization of NP-KLH and found no differences in the frequency of cells at different phases of cell cycle (Fig. S4O-R). These data suggest that STAT3 deficiency minimally affects GC B cell proliferation which may not fully account for increased LZ and reduced DZ of STAT3 deficient GCs.

### STAT3 deficiency reduces recycling and long-lived plasma cell numbers but increases memory B cells

To determine how STAT3 may control the GC output by regulating the DZ and LZ organization, we employed the GC B cell fractionation approach (Inoue et al., 2021; Ise et al., 2018). According to this method, GC B cells can be categorized into 7 different fractions based on CD38, CD69, Bcl-6, and Ephrin b1 (Efnb1) expression, where each fraction is associated with a specific B cell fate (Fig. 4A, D). CD38^-^ LZ GC B cells can be categorized into four Fractions (Fig. 4A) based on CD69 and Bcl-6 expression in which Fraction 1 (CD38^-^Bcl^-^ 6^lo^CD69^hi^) and Fraction 2 (CD38^-^Bcl^-^6^hi^CD69^hi^) cells represent precursors of plasmablasts and recycling GC cells, respectively (Ise et al., 2018). Fraction 3 (CD38^-^Bcl^-^6^hi^CD69^lo^) demarcates pre-fate LZ cells awaiting selection signals and Fraction 4 (CD38^-^Bcl^-^6^lo^CD69^lo^) contains memory precursors (Ise et al., 2018). Similarly, NP^+^ B cells transitioning towards memory re-express CD38 and can be separated into three Fractions (Fig. 4D) based on Bcl-6 and Efnb1 expression in which Fraction 5 (CD38^+^Bcl^-^6^+^Efnb1^+^), Fraction 6 (CD38^+^Bcl^-^6^lo^Efnb1^+^) and Fraction 7 (CD38^+^Bcl^-^6^-^Efnb1^-^) represent pro-, pre-, and mature-memory B cells, respectively (Inoue et al., 2021). We found significantly reduced percentages and numbers of recycling cells (Fraction 2) within the GC LZ, whereas memory precursors (Fraction 4) and mature-memory (Fraction 7) cells were not reduced, but rather were increased in STAT3^fl/fl^CD23^Cre^ mice compared to CD23^Cre^ control mice on 14d post-NP-KLH-immunization (Fig. 4A-F).

**Fig. 4.**
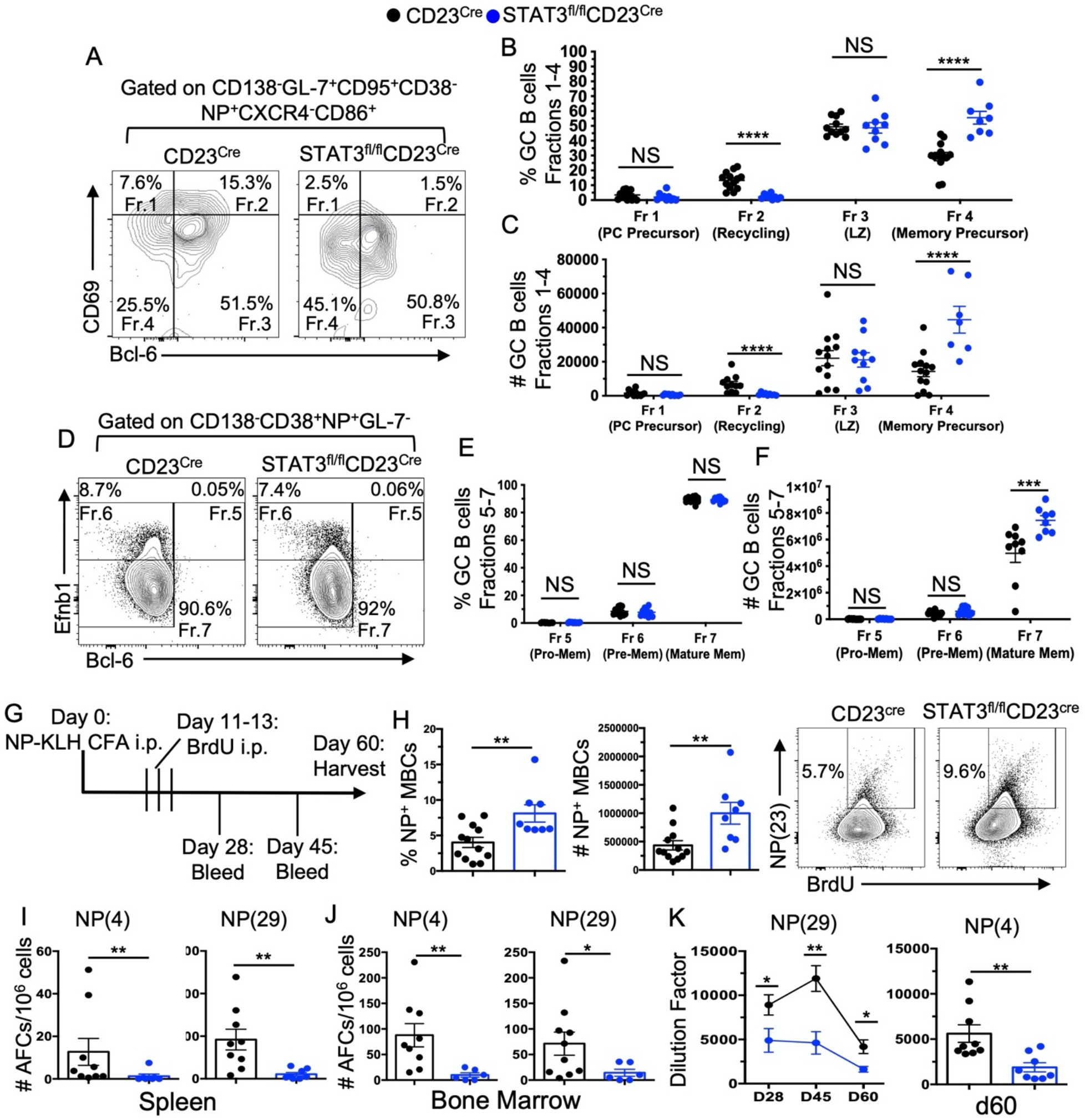
B cell intrinsic STAT3 signaling is required for GC B cell recycling and long-lived plasma cell differentiation but not memory formation. **(A-F)** Flow cytometric examination of GC B cell fractions 14d post-immunization in CD23^Cre^ and STAT3^fl/fl^CD23^Cre^ mice. (**A**) Gating strategy, (**B**) percentage and (**C**) number of GC B cell fractions 1-4 associated with LZ GC-fate pre-gated on CD138^-^B220^+^GL-7^+^CD95^+^CD38^-^NP^+^CXCR4^-^CD86^+^ cells. (**D**) Gating strategy, (**E**) percentage and (**F**) number of GC B cell fractions 5-7 associated with post-GC memory committed cells pre-gated on CD138^-^CD38^+^NP^+^GL-7^-^. (**G**) Schematic of immunization and BrdU pulsing used for assessment of memory formation in CD23^Cre^ and STAT3^fl/fl^CD23^Cre^ mice. **(H**) Percentage, number, and representative flow plot of B220^+^CD38^+^CD95^-^CD138^-^NP^+^BrdU^+^ MBCs on 60d post-NP-KLH immunization in CD23^Cre^ and STAT3^fl/fl^CD23^Cre^ mice. Number of **(I)** splenic and (**J**) bone marrow NP4- and NP29-specific IgG1 producing AFCs on 60d post-NP-KLH immunization. **(K)** Quantification of NP29- and NP4-specific IgG1 serum titers at indicated time points. Each symbol represents an individual mouse and data are presented as means ±SEM (n=6-14 mice per group). Data in each panel represent two-three experiments. P values were calculated via an unpaired t-test or Mann-Whitney test (NS, p>0.05, *, p <0.05, **, p <0.01, ***, p <0.001, ****).

To confirm the role of STAT3 signaling in B cells in long term memory formation, STAT3^fl/fl^CD23^Cre^ and CD23^Cre^ mice were immunized with NP-KLH followed by an injection of BrdU (i.p.) every 12 hours on d11-13 of the GC response (Fig. 4G), as previously described (Weisel et al., 2016). B220^+^NP^+^BrdU^+^ memory B cells (MBCs) were analyzed by flow cytometry 60d post-immunization. STAT3^fl/fl^CD23^Cre^ mice had an increased percentage and number of MBCs compared to CD23^Cre^ control mice (Fig. 4H). However, numbers of high affinity (NP4) and total (NP29) long-lived antibody-forming cells (AFCs) at the d60 time point were significantly lower in the spleen and BM of STAT3^fl/fl^CD23^Cre^ mice (Fig. 4I, J). In line with reduced long-lived AFC responses, NP4- and NP29-specific prolonged (d28, d45 and d60) IgG1 Ab responses were also diminished in STAT3^fl/fl^CD23^Cre^ mice compared to CD23^Cre^ control mice (Fig. 4K). Together, our data show significantly reduced numbers of recycling LZ cells (Fraction 2) and GC-derived LL-PCs in the absence of STAT3 and that STAT3 deletion increases MBC number.

### STAT3 regulates LZ B cell number and recycling into the DZ but not GC initiation or maintenance

To confirm the B cell intrinsic role of STAT3 in regulating LZ B cell recycling into the DZ, we utilized a tamoxifen inducible B cell-specific STAT3 deletion system by crossing STAT3^fl/fl^ mice with hCD20^ERT2-Cre^ mice (Khalil et al., 2012). We first treated STAT3^fl/fl^hCD20^ERT2-Cre^ and hCD20^ERT2-Cre^ mice with tamoxifen on d0-d5 post-immunization (Fig. 5A) at the GC initiation phase and assessed DZ and LZ distribution on d7 when GCs become polarized (MacLennan, 1994; Victora and Nussenzweig, 2012). Total and NP-specific GC responses were intact in STAT3^fl/fl^hCD20^ERT2-Cre^ mice compared to hCD20^ERT2-Cre^ control mice (Fig. 5B, C). Notably, STAT3 deletion in B cells resulted in accumulation of B cells in the LZ as evidenced by significantly increased LZ and decreased DZ B cells in STAT3^fl/fl^hCD20^ERT2-Cre^ mice compared to hCD20^ERT2-Cre^ control mice (Fig. 5D, E). Percentages of NP-specific PCs were reduced in STAT3 deleted mice (Fig. 5F, G). We then treated STAT3^fl/fl^hCD20^ERT2-Cre^ and hCD20^ERT2-Cre^ control mice with tamoxifen during the peak GC response (on d11-13) post-NP-KLH immunization, when GC B cells are fully organized into the DZ and LZ of the GC, prior to analyzing these mice on d14 (Fig. 5H, I). Total GC B cell responses in STAT3^fl/fl^hCD20^ERT2-Cre^ mice were higher than in hCD20^ERT2-Cre^ mice, although NP-specific GC responses showed no difference (Fig. 5J). Similar to d7 data, we observed a reduced frequency of DZ and increased frequency of LZ B cells in STAT3^fl/fl^hCD20^ERT2-Cre^ mice compared to control mice (Fig. 5K, L).

**Fig. 5.**
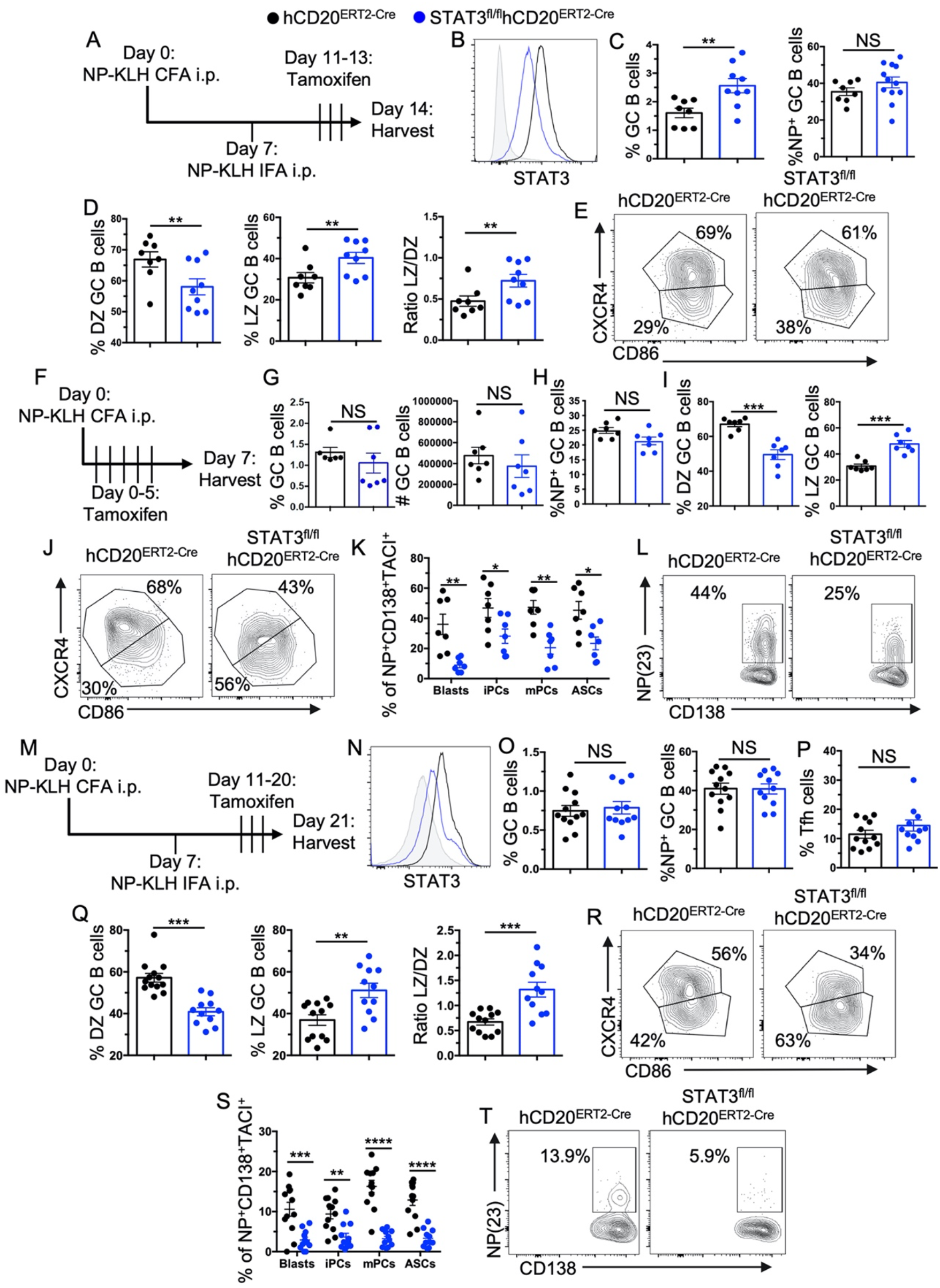
STAT3 signaling in B cells regulates LZ B cell recycling into the DZ but not GC initiation or GC B cell maintenance. **(A-E)** Examination of B cell intrinsic STAT3 deletion after GC establishment. **(A)** Schematic of NP-KLH immunization and Tamoxifen administration schedule in hCD20^ERT2-Cre^ and STAT3^fl/fl^hCD20^ERT2-Cre^ mice. **(B)** Representative histograms of STAT3 expression in hCD20^ERT2-Cre^ (black) and STAT3^fl/fl^hCD20^ERT2-Cre^ (blue) mice relative to an isotype control (gray). **(C)** Percentage of B220^+^GL-7^+^CD95^+^ GC B cells from indicated mice at 14d post-immunization. **(D)** Percentage of DZ and LZ GC B cells and the ratio of LZ/DZ GC B cells from these mice. **(E)** Representative flow plots of LZ and DZ GC B cells. **(F-L)** Examination of B cell intrinsic STAT3 deletion during GC initiation. **(F)** Schematic of NP-KLH immunization and Tamoxifen administration schedule in hCD20^ERT2-Cre^ and STAT3^fl/fl^hCD20^ERT2-Cre^ mice. **(G)** Frequency and number of splenic GC B cells (B220^+^GL-7^+^CD95^+^). **(H)** Percentage of NP+ GC B cells from indicated mice. **(I)** Percentages of DZ (CXCR4^+^CD86^-^) and LZ (CXCR4^-^CD86^+^) GC B cells on d7 post-NP-KLH immunization. **(J)** Representative flow plots depicting LZ/DZ distribution. **(K)** Frequency of IgD^-^NP^+^CD138^+^TACI^+^ ASCs and subpopulations of plasma cells as described in the legend to Figure 1. **(L)** Representative flow plot of IgD^-^NP^+^CD138^+^TACI^+^ plasma cells. **(M-T)** GC B cell responses in extended STAT3 deletion post-GC establishment. **(M)** Schematic of NP-KLH immunization and Tamoxifen administration schedule in hCD20^ERT2-Cre^ and STAT3^fl/fl^hCD20^ERT2-Cre^ mice. **(N)** Representative histograms of STAT3 expression in hCD20^ERT2-cre^ (black) and STAT3^fl/fl^hCD20^ERT2-cre^ (blue) mice relative to an isotype control (gray). **(O)** Percentage of B220^+^GL-7^+^CD95^+^ GC B cells 21d post-immunization. **(P)** Percentage of CD4^+^CD44^+^CD62L^-^PD-1^+^CXCR5^+^ Tfh. **(Q)** Percentage of DZ and LZ and ratio of LZ/DZ GC B cell populations from indicated mice. **(R)** Representative flow plots of LZ and DZ B cells. **(S)** Percentages of NP-specific plasma cell populations as described in the legend to Figure 1. (**T**) Representative flow plot of IgD^-^NP^+^CD138^+^TACI^+^ plasma cells. Each symbol represents an individual mouse and data are presented as means ± SEM. Data in each panel represent two-three experiments. P values were calculated via an unpaired t-test or Mann-Whitney test (NS, p>0.05, *, p<0.05, **, p <0.01, ***, p <0.001, ****, p <0.0001).

The role of STAT3 in GC maintenance was previously described using STAT3^fl/fl^CD19^Cre^ mice (Ding et al., 2016) in which STAT3 was constitutively deleted during the pro-B cell stage of B cell development. Here we evaluated the definitive B cell-intrinsic role of STAT3 in GC maintenance by treating STAT3^fl/fl^hCD20^ERT2-Cre^ and hCD20^ERT2-Cre^ mice with tamoxifen on d11-d20 (during the GC maintenance stage) post-NP-KLH immunization, then analyzed these mice on d21 (Fig. 5M, N). We found no differences in total and NP-specific GC (Fig. 5O) and Tfh (Fig. 5P) responses in mice with STAT3 deletion in B cells compared to treated hCD20^ERT2-Cre^ control mice. Consistently, we observed a reduced frequency of DZ B cells and an increased frequency of LZ B cells in STAT3^fl/fl^hCD20^ERT2-Cre^ mice than in hCD20^ERT2-Cre^ control mice (Fig 5Q, R). We also observed reduced percentages of various subsets of total and NP-specific plasma cells (Fig. 5S, T). These data together with the GC fractionation data (Fig. 4A-F) highlight the B cell-specific role of STAT3 in regulating LZ B cell number and recycling into the DZ, but not in the GC initiation or maintenance.

### Th cell-dependent activation of STAT3 in LZ B cells regulates their recycling

To delineate the mechanisms by which STAT3 signaling in B cells regulates LZ B cell recycling into the DZ, we measured total STAT3 expression in DZ and LZ B cells on d14 post-immunization of C57BL/6 (B6) mice and found no significant difference in expression between the zones (Fig. 6A). Maximal transcriptional activity of STAT3 is mediated by dual phosphorylation of tyrosine 705 (STAT3 pY705) and serine 727 (STAT3 pS727) (Wen et al., 1995). We found a significantly higher percentage of pY705^+^ and pS727^+^ GC B cells in the LZ than in the DZ (Fig. 6B). T cell help is critical for selection of LZ B cells into recycling GC cells (Finkin et al., 2019; Ise et al., 2018; Victora et al., 2010). We hypothesized that a T-dependent signal promotes STAT3 activity in LZ B cells to drive their recycling into the DZ. We treated B6 mice with the MR1 monoclonal antibody, which blocks CD40L-CD40 interactions between Tfh and GC B cells, on d13.5 post-immunization prior to analyzing on d14. MR1-treated mice had a significantly reduced percentage of phosphorylated STAT3 Y705 and S727 LZ B cells than control mice, whereas no difference was observed in DZ B cells (Fig. 6C, D).

**Fig. 6.**
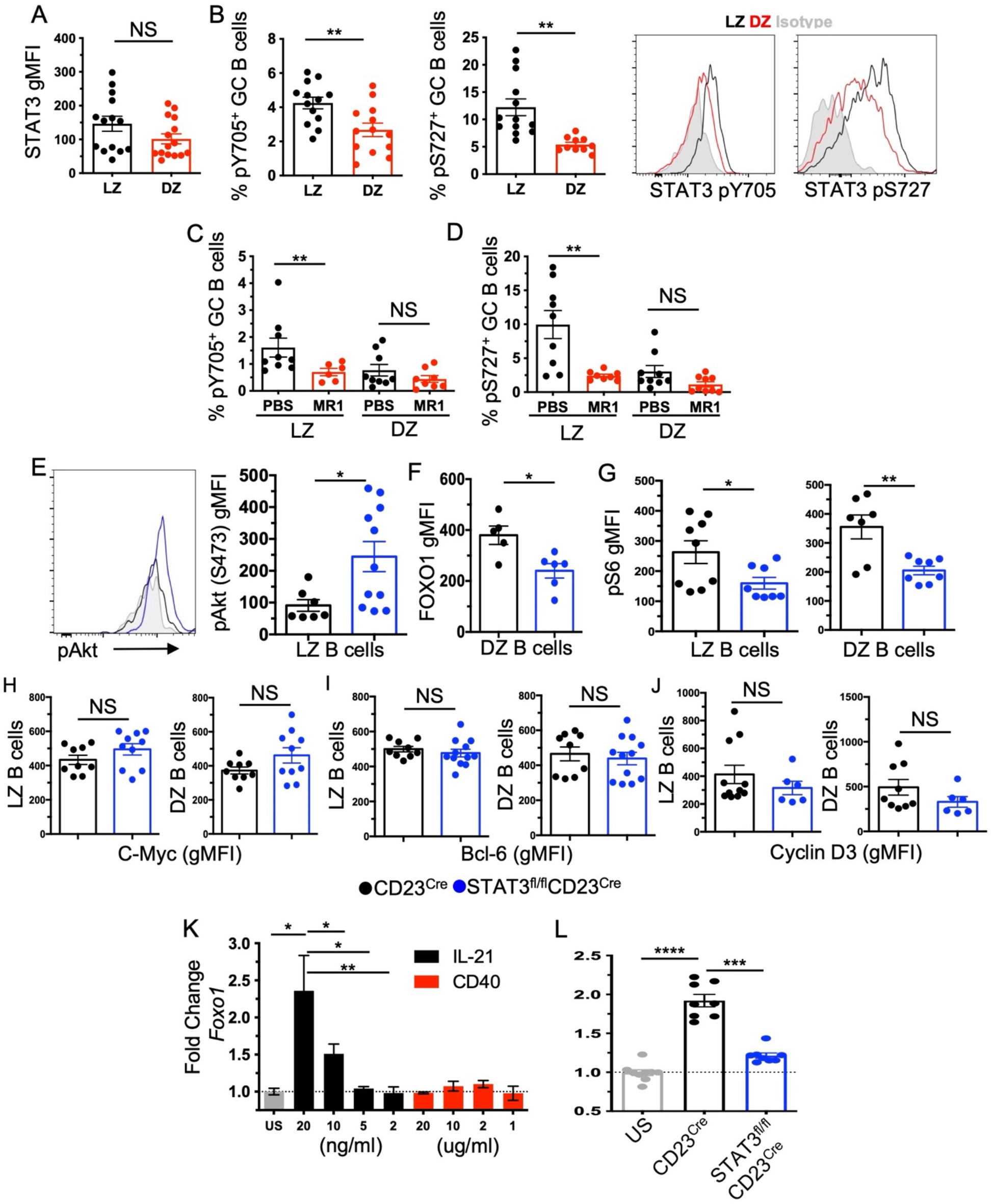
T-dependent LZ signaling promotes STAT3 activity and signaling associated with DZ recycling. **(A)** Flow cytometric analysis of STAT3 expression (gMFI) in LZ and DZ compartments of B220^+^GL-7^+^CD95^+^ GC B cells on14d post-NP-KLH immunization of B6 mice. (**B**) Percentage of STAT3 pY705^+^ and pS727^+^ GC B cells within each zone and representative histograms (LZ: black, DZ: blue). (**C, D**) Day 13.5 post-immunization B6 mice were injected (i.v.) with anti-CD154 (clone MR-1) and the percentage of phosphorylated Y705 and S727 GC B cells within each zone measured. (**E-J**) Examination of signaling proteins associated with DZ recycling 14d post-immunization in CD23^Cre^ and STAT3^fl/fl^CD23^Cre^ mice. (**E**) Representative histogram of endogenous pAKT (Ser473) and quantification of pAKT (gMFI) in LZ GC B cells. (**F**) FOXO1 expression (gMFI) in DZ GC B cells. (**G**) Phosphorylated S6 (Ser 235/236) in LZ and DZ GC B cells. Expression (gMFI) of cMyc (**H**), Bcl-6 (**I**), and Cyclin D3 (**J**) in LZ and DZ B cells from indicated mice. (**K**) *In vitro* generated GC B cells (iGC) from B6 mice were stimulated with the indicated amount of IL-21 (ng/mL) or anti-CD40 (µg/mL) for 12 hours and *Foxo1* levels were compared relative to β-actin by RT-PCR. (**L**) *Foxo1* levels were measured by RT-PCR in iGC B cells generated from CD23^Cre^ and STAT3^fl/fl^CD23^Cre^ mice and stimulated for 12 hours with 20ng/mL of IL-21. Each symbol represents an individual mouse (n=6-12 mice per group) and data are presented as means ±SEM. Data in each panel represent three-four experiments. P values were calculated via an unpaired t-test, Mann-Whitney test, or an ANOVA (NS, p>0.05, *, p <0.05, **, p <0.01, ***, p <0.001, ****, p <0.0001).

Based on the confirmation of STAT3 activation in LZ B cells by T cell derived signals, we investigated the impact of STAT3 deficiency on signals induced by Tfh-selection. GC cell recycling from the LZ to DZ was shown to be mediated by reduction in pAkt, and increased expression/activity of FOXO1, Myc, and mTOR/pS6 (Dominguez-Sola et al., 2015; Dominguez-Sola et al., 2012; Ersching et al., 2017; Finkin et al., 2019; Inoue et al., 2017; Luo et al., 2018; Roberto et al., 2021; Sander et al., 2015). Compared to control mice, STAT3^fl/fl^CD23^Cre^ mice had elevated levels of pAKT in LZ B cells (Fig 6E), paired with reductions in DZ expression of FOXO1 (Fig. 6F) on 14d post-immunization. Further, we observed reductions in pS6 in both LZ and DZ compartments relative to control mice (Fig. 6G). We did not find differences in Bcl-6, Myc, or Cyclin D3 expression (Fig. 6H-J). FOXO1 is a STAT3 target in T cells (Oh et al., 2011) but has not been explored in GC B cells. Given the established role of FOXO1 in regulating DZ, we delineated the role of STAT3 in the induction of FOXO1 by utilizing a well-established 40LB GC culture system (Nojima et al., 2011). We found a dose-dependent increase in *Foxo1* expression following IL-21 but not CD40 stimulation (Fig. 6K) that was dependent on STAT3 expression (Fig. 6L). Together, our data demonstrate that T-dependent signals mediate STAT3 activation in LZ B cells necessary for LZ B cell recycling into the DZ.

### STAT3 signaling in B cells controls GC zone organization by regulating and targeting genes required for DZ recycling

To comprehensively define the role of STAT3 in regulating the GC B cell transcriptome, we sorted GC B cells from STAT3^fl/fl^CD23^Cre^ and CD23^Cre^ control mice on d14 post-NP-KLH immunization and performed RNAseq. We identified 1,086 genes differentially expressed (367 downregulated and 719 upregulated) in STAT3^fl/fl^CD23^Cre^ GC B cells (Figs. 7A, S5A). Several of these differentially expressed genes were previously shown to regulate GC responses including *Cxcr4* (CXCR4), *Ccl22* (CCL22), *Tet2* (TET2), *Aicda* (AID), *Tox2* (TOX2), *Fcer2a* (CD23), and others (Bannard et al., 2013; Dominguez et al., 2018; Liu et al., 2021; Mesin et al., 2016; Nakagawa et al., 2021; Xu et al., 2019). GSEA transcriptionally confirmed the alterations to GC DZ and LZ compartmentalization in the absence of STAT3 (Figs. 7B, S5B). We also identified numerous T cell-driven transcriptional programs in GC B cells critical for DZ recycling, including Myc and E2F targets, and mTORC1 signaling which were enriched in controls relative to STAT3-deficient GC B cells (Fig. 7B, S5B). These data suggest that STAT3 deficiency results in alterations to the transcriptome of GC B cells resulting in defects in the GC organization of DZ and LZ in a T cell dependent fashion.

**Fig. 7.**
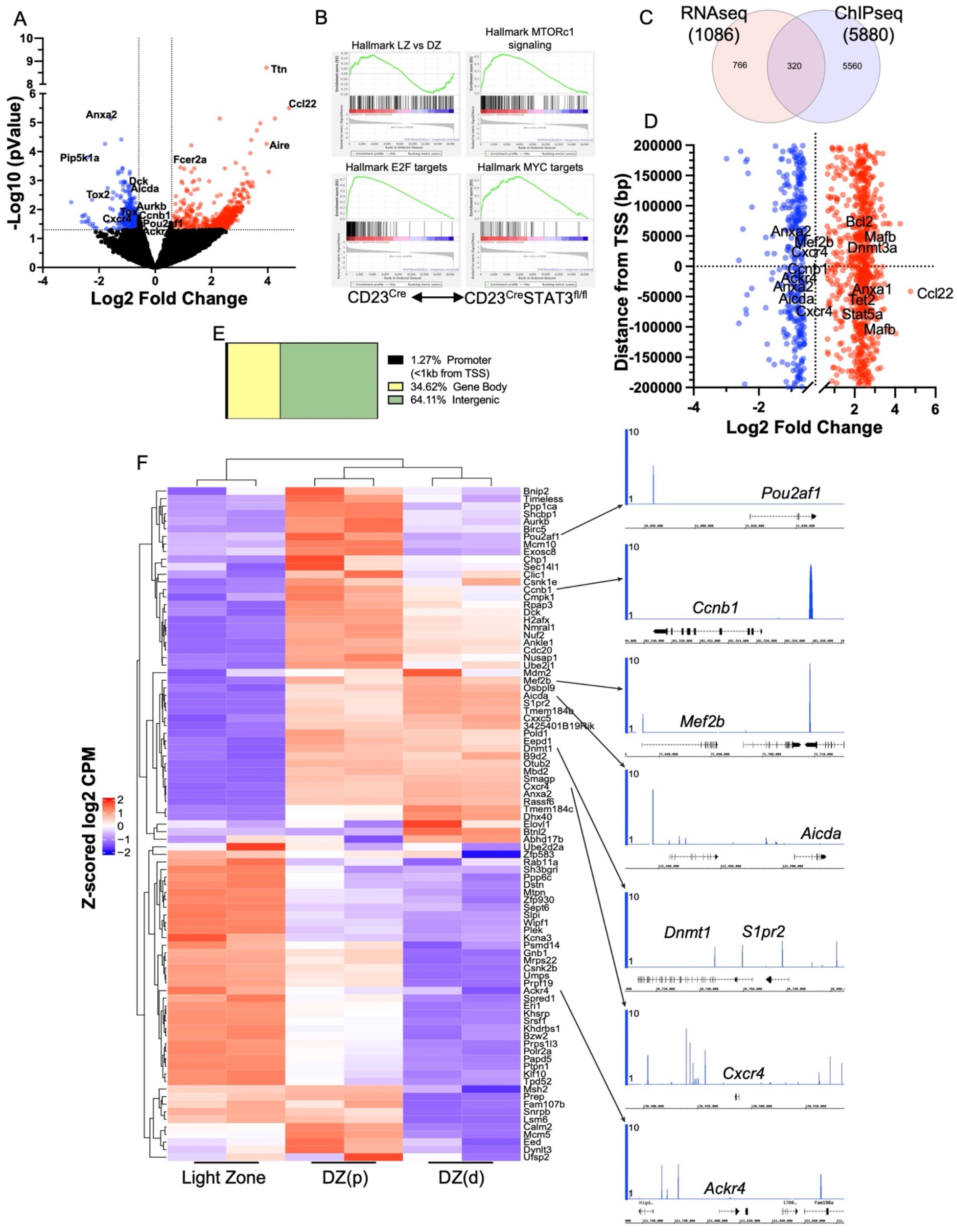
STAT3 signaling in B cells controls the GC organization by targeting genes required for DZ recycling. (**A**) RNAseq analysis of the differentially expressed genes identified in GC B cells from STAT3^fl/fl^CD23^cre^ and CD23^cre^ mice on 14d post-NP-KLH immunization. (**B**) Gene set enrichment analysis (GSEA) depicting gene enrichment in hallmark LZ vs DZ gene signatures, mTORC1 signaling, E2F targets, and Myc targets. (**C**) Venn diagram depicting the overlap of genes identified by RNAseq (logFC > 1.5, FDR > 0.05) and ChIPseq. (**D**) Plot of differentially expressed genes identified from RNAseq versus the distance of a STAT3 peak to the gene’s TSS. **(E)** Genomic locations of STAT3 ChIPseq peaks using HOMER. **(F)** RNAseq heatmap of genes upregulated in CD23^Cre^ versus STAT3^fl/fl^CD23^Cre^ GC B cells with STAT3 binding within 50 kb of the TSS. Heatmap shows RNAseq expression across the light zone (LZ), dark zone proliferation (DZp), and dark zone differentiation (DZd). STAT3 ChIPseq tracks at genes of interest. Tracks are representative of duplicate ChIPseq samples.

To identify the genes directly regulated by STAT3, we performed ChIPseq on GC B cells sorted from B6 mice on d14 post-NP-KLH immunization. We identified 11,108 STAT3 peaks mapping to 5,880 genes, 320 of which were also identified in our RNAseq analysis (Fig. 7C, D). The majority of the STAT3 peaks were found within intergenic regions (64.1%), followed by gene bodies (34.6%), and a small percentage (1.3%) at the promoter (<1kb from the transcription start site, TSS) (Fig. 7E). We mapped the distance between the TSS of differentially expressed genes identified by RNAseq and the closest STAT3 peaks identified by ChIPseq (Fig. 7F). Consistent with our RNAseq data, we identified that STAT3 bound nearby several key genes related to DZ recycling and GC biology, including *Cxcr4, Aicda, Ccl22, Ccnb1*, and others (Figure 7F) (Bannard et al., 2013; Kennedy et al., 2020; Liu et al., 2021; Victora and Nussenzweig, 2012; Young and Brink, 2021).

The DZ has recently been divided into two distinct niches, one in which GC B cells proliferate (DZp) and the other where they differentiate (DZd), increase AID activity, and undergo SHM of the BCR (Kennedy and Clark, 2021; Kennedy et al., 2020). To identify the role of STAT3 in regulating the transcriptional program that develops as T cell-selected GC B cells transition from the LZ to the DZp and then to the DZd, we first identified STAT3 peaks within 50kb of our differentially expressed genes’ TSS. We then determined the expression profile of these genes across the LZ, DZp, and DZd zones using previously described RNAseq data (Kennedy et al., 2020) (Fig. 7F, left panel). We identified that STAT3 bound nearby numerous genes that were differentially expressed as LZ GC B cells transited through the DZp and DZd (Fig. 7F, right panel). These include *Pou2af1, Ccnb1, Mef2b, Aicda, Dnmt1, S1pr2, Cxcr4*, and *Ackr4*. Consistent with our data, CXCR4-deficient cells were previously shown to be restricted in the LZ (Bannard et al., 2013). These data highlight a previously unrecognized role for B cell intrinsic STAT3 in the transcriptional regulation of GC B cells recycling out of the LZ into the DZ.

## DISCUSSION

GC B cell and Tfh collaboration in the GC LZ is a major checkpoint that determines whether GC B cells egress the GC as plasma or memory B cells, reenter the DZ for further proliferation and SHM, or undergo apoptosis (Ise et al., 2018; Mesin et al., 2016; Shinnakasu and Kurosaki, 2017; Shlomchik et al., 2019; Weisel et al., 2016). Discrete organization of the GC zones is believed to be important for optimal GC responses and affinity maturation (Bannard and Cyster, 2017; Laidlaw and Cyster, 2021; Young and Brink, 2021), but the overlap between the T-dependent signals in the LZ and the GC B cell fate decision is incompletely understood. Here, we identified a previously unappreciated role for STAT3 in the regulation of GC B cell zonal organization. We provided evidence that T-cell dependent STAT3 signaling in LZ B cells controls their number and recycling into the DZ and the maturation of GC-derived LL-PCs, and that STAT3 deficiency favors the differentiation of MBCs. This functionality is mediated by STAT3 targeting genes required for LZ B cells to recycle and transit through the proliferation and differentiation stages of the DZ. We further discovered that organization of the GC zones is not necessary for overall GC response, GC maintenance, or SHM of Ig genes.

Several previous studies described the B cell intrinsic role of STAT3 in the regulation of PC and GC responses (Dascani et al., 2018; Ding et al., 2016; Fornek et al., 2006; Kane et al., 2016). While these studies agreed on the crucial role of STAT3 in PC differentiation and IgG Ab responses, conflicting findings on the GC responses were reported. Using STAT3^fl/fl^CD19^cre^ mice, Fornek et al. originally identified no impact of STAT3 deficiency on the GC response (Fornek et al., 2006), whereas other groups found reduced GC responses in these mice (Dascani et al., 2018; Ding et al., 2016). A reduced GC response was also recapitulated in a transfer model in which B cells from CD19^cre^STAT3^fl/fl^ mice expressing a transgenic BCR specific for hen egg lysozyme (SW_HEL_) were transferred to wild type recipients (Kane et al., 2016). However, detailed mechanisms of STAT3 functions in GC regulation and GC output were not defined. To delineate such mechanisms, we employed a prime-boost immunization approach to induce a robust GC reaction. Similar to a previous study (Fornek et al., 2006), we found no effect of STAT3 on overall GC responses using this approach in three B cell conditional KO mice and one inducible B cell KO system. Despite a quantitatively normal GC response, DZ and LZ compartmentalization was altered in the absence of STAT3 in B cells.

Cell death and proliferation must be balanced to maintain the GC response (Kennedy and Clark, 2021; Mesin et al., 2016). GC B cell proliferation is anatomically separated, where 10-15% of LZ cells are thought to proliferate while the majority of proliferation occurs in the DZ (Shlomchik et al., 2019). Apoptosis can occur in both zones through distinct zone-specific mechanisms. While previous studies attributed reduced GC responses to increased apoptosis of GC B cells deficient in STAT3 (Ding et al., 2016; Kane et al., 2016), we found no major effect of STAT3 deficiency on GC B cell proliferation or apoptosis in either zone. This was not surprising given STAT3 deficiency had no impact on total GC response in our case. We favor the possibility that discrepancies in GC responses and rate of GC B cell apoptosis were due to the robustness of the GC responses induced in various studies.

During the preparation of our current manuscript two studies reported a B cell intrinsic role for IL-21 signaling in GC B cell proliferation as a cause of GC DZ and LZ disorganization in IL-21R^-/-^ mice (Dvorscek et al., 2022; Zotos et al., 2021). We, however, observed no significant deficit in overall B cell proliferation during initiation and peak GC responses, although we observed a modest reduction in the post-S phase LZ GC B cells in the absence of STAT3. We also found that STAT3 deficiency modestly affected LZ GC B cell progression through S phage of the cell cycle. These results fit with recent findings of B cell intrinsic IL-21R signaling in promoting LZ B cell entry into the S phase of the cell cycle (Zotos et al., 2021). However, earlier, and recent papers examining the GC response in IL-21R^-/-^ mice demonstrated that these mice failed to elicit sustained GC responses beyond day 7 post-immunization (Linterman et al., 2010; Quast et al., 2022; Zotos et al., 2010; Zotos et al., 2021); thus, defects in proliferation are more likely to affect the GC response and GC DZ/LZ distribution. In contrast, overall GC responses in STAT3 deficient animals were comparable to wild-type controls and persisted at least until d21, despite altered zonal organization. These data suggest that IL-21 signaling in B cells controls GC zone distribution and GC output through STAT3-dependent and -independent mechanisms in which STAT3-independent mechanism provides the survival and proliferation signals to B cells. Our current data delineate the mechanisms by which STAT3 regulates GC DZ/LZ distribution and GC output of long-lived plasma cells and memory B cells.

STAT3 is activated by several stimuli that have been implicated in GC biology. While our data suggest a link between IL-21 signaling, STAT3, and the induction of FOXO1, as described above it is unlikely that IL-21 signaling is the exclusive stimulus regulating STAT3 mediated LZ and DZ organization. A previous study demonstrated that loss of IL-10 production by Tfh cells resulted in reduced DZ and increased LZ B cells without GC collapse (Laidlaw et al., 2017).

Although similar to our findings, the DZ/LZ phenotypes in our STAT3-deficient mice were more dramatic than those observed in the study by Laidlaw et al (Laidlaw et al., 2017). Other STAT3 activating cytokines such as Type I interferon, IL-6, IL-2, and IL-7 have also been implicated in various aspects of GC formation or maintenance (Arkatkar et al., 2017; Ballesteros-Tato et al., 2012; Ersching et al., 2017; Hanissian and Geha, 1997; Laidlaw et al., 2017; Ray et al., 2014; Tangye and Ma, 2020; Xu et al., 2017), but to our knowledge no effects of these cytokines on GC zonal organization have been reported. In agreement with the alterations to the LZ and DZ organization, we identified reductions in FOXO1 expression and phosphorylated S6 (downstream of mTORC1) in DZ cells, paired with increases in phosphorylated Akt in LZ cells in the absence of STAT3. Of note, we detected no significant differences in the expression of Myc or Cyclin D3 either in DZ or LZ cells, although the methods we used may not provide the resolution to detect the small percentage of cells upregulating these molecules. Thus, we propose that STAT3 activation downstream of multiple T-dependent stimuli provides the initial signal required to poise Tfh-selected GC B cells to recycle towards the DZ.

FOXO1 is a known regulator of GC DZ and is induced by a T-dependent signal (Dominguez-Sola et al., 2015; Inoue et al., 2017; Roberto et al., 2021; Sander et al., 2015). Our data suggest a role for STAT3 in the regulation of FOXO1 in GC B cells, although we did not identify *Foxo1* as a target gene of STAT3 in our ChIP-seq analysis. This could be due to an insufficient signal for detecting STAT3 binding to the *Foxo1* gene using the large-scale approach of ChIP-seq in total GC B cells or that STAT3 may cooperatively regulate *Foxo1* through interactions with other transcription factors. An alternative scenario is that STAT3 is targeting a regulatory element such as an enhancer to support the induction of FOXO1 expression. Given the role of FOXO1 in DZ regulation, it was surprising that alteration to GC DZ organization in the absence of FOXO1 had no major effect on SHM analyzed on the VH186.2 family at 14d post-immunization, although affinity maturation was affected (Dominguez-Sola et al., 2015; Sander et al., 2015). In line with this, we also found no major impact of GC zone disorganization mediated by STAT3 deficiency, on SHM and affinity maturation through VH186.2 cloning and sequencing at d14 post-immunization or through BCR repertoire sequencing of NP^+^ and NP^-^ GC B cells at d21 post-immunization. Collectively, published and our data together suggest that SHM is a dynamic process which may require only a partially functional DZ.

We observed an accumulation of B cells in the LZ in the absence of STAT3 which was associated with an increase in MBCs. BCR stimulation paired with weak T cell help was previously shown to promote GC B cell exit as MBCs (Inoue et al., 2021; Ise et al., 2018; Laidlaw and Cyster, 2021; Shinnakasu and Kurosaki, 2017). Our data indicate that STAT3 functions as a negative regulator of GC B cells exit as MBCs. Consistent with these data we identified numerous STAT3 binding sites near genes *Tle3, Klf2*, and *Zeb2* that have been shown to play important roles in MBC development (Laidlaw et al., 2020). The other possibility is that that the increase in GC-derived MBCs is the result of an increased number of LZ B cells receiving a relatively low level of T cell help due to the loss of STAT3 signaling in GC B cells, thus promoting their exit from the GC as MBCs. Future studies can more precisely focus on interrogating the relationship between STAT3, the generation of GC derived memory, and STAT3-dependent signals that mediate the MBC phenotype.

STAT3 has a known role in the differentiation of plasma cells. Previous work examining the effect of STAT3 haploinsufficiency on GC and PC fate of transferred B1-8^hi^ cells (Ise et al., 2018) and our current study of STAT3 deficiency in B cells on polyclonal GC and PC responses have demonstrated that STAT3 signaling in GC B cells is not required for the initial commitment of GC B cells to a plasma cell fate, but rather is necessary for post-GC PC differentiation. We found that STAT3 was involved in the formation of antigen-specific splenic plasmablasts and long-lived bone marrow PCs/AFCs, which agree with findings in human patients with STAT3 loss-of-function mutations (Avery et al., 2010; van de Veen et al., 2019). Our data indicate that only after GC cells are committed to become plasma cells and begin to express BLIMP1, does STAT3 seem to be involved. Future studies will be required to determine the temporal relationship between IRF4 and STAT3 in the induction of BLIMP1 for plasma cell development.

Through transcriptional analysis we identified that STAT3 is involved in the regulation of multiple genes in GC B cells with known roles in LZ cell recycling and transiting through proliferation and differentiation stages of DZ B cells (Kennedy and Clark, 2021; Kennedy et al., 2020). Although there were limited STAT3 binding peaks located directly at the TSS of genes, we found that most STAT3 peaks were located adjacent to or near genes critical for the GC response. Given the differential expression profile, our data suggest that STAT3 is involved in regulating these genes by binding regulatory elements such as distal promoters or enhancers thereby fine-tuning the GC reaction. Specifically, we identified numerous STAT3 binding sites near *Cxcr4* and *Ccnb1*, both of which were downregulated in STAT3 deficient GC B cells and are known to be involved in DZ migration and cell cycle progression, respectively (Bannard et al., 2013; Kennedy et al., 2020). Further, we identified STAT3 involvement in the regulation of *Aicda, Pou2af1, Mef2b, Dnmt1*, and *S1pr2* all of which are upregulated as GC B cells transition from the LZ to the DZp (Kennedy et al., 2020). Given that the RNAseq was performed on total GC B cells, enrichment of subpopulations such as recently selected LZ GC B cells, may reveal a higher resolution to identify the transcriptional differences mediated by STAT3.

A limitation to our studies is that we focused on STAT3 transcriptional control of the GC response and GC output; however, STAT3 has known non-transcriptional activities that could be involved in GC biology. Notably, stimulation of mTORC1 leads to phosphorylation of STAT3 Serine 727 allowing its entry into the mitochondria to support oxidative phosphorylation (Bernier et al., 2011; Kim et al., 2009; Lee et al., 2015; Meier and Larner, 2014; Wegrzyn et al., 2009; Yokogami et al., 2000). Given the induction of mTORC1 upon GC B cell selection (Ersching et al., 2017) and our findings, it is interesting to speculate that STAT3 may contribute to the metabolic programming required for DZ proliferation. Future studies will be necessary to parse out the transcriptional and non-transcriptional functions of STAT3 in GC regulation.

During a pathogenic infection fine-tuning GC B cell selection by Tfh cells is a critical step for the development of high affinity antibody producing LL-PCs and MBCs. Our data demonstrate that STAT3 is a key player regulating this GC checkpoint. Identifying signals involved in this critical B cell developmental checkpoint, and the relationship between and functions of various zone-specific signals is integral to improving our understanding of GC biology. A comprehensive understanding of GC biology will likely aid in the development of new therapeutics or improved vaccines.

## MATERIALS AND METHODS

### Study Design

The overall goal of this study was to delineate the mechanisms by which B cell intrinsic STAT3 regulates the GC response and GC output of long-lived plasma and memory B cells. This was accomplished by using several developmental stage and tissue specific B cell STAT3 conditional and temporally controlled tamoxifen inducible knockout systems. The phenotypic examination of various B and T cell populations and expression of signaling molecules in GC B cells were performed by flow cytometry and immunofluorescence microscopy. Further mechanistic studies were performed through *in vivo* treatments of mice, the generation of *in vitro* differentiated GC B cells, and the analysis of RNAseq, ChIPseq and BCR sequencing data. The number of mice per experiment was determined based on the power calculation of our published data (Chodisetti et al., 2020a; Domeier et al., 2018; Fike et al., 2021). Littermate controls were used where possible. Control and experimental mice were age- and sex-matched. All data points generated in various experiments were included in the analysis. All experimental findings were replicated and the number of replicates are indicated in the figure legends.

### Mice

C57BL/6 (B6), B6.129S1-*Stat3*^*tm1Xyfu*/^J (STAT3^flox^), B6.129P2(C)-*Cd19*^*tm1(cre)Cgn*^*/*J (CD19^cre^), B6.Cg-Tg(Cd4-cre)1Cwi/Bfluj (CD4^cre^), and B6.129P2(Cg)-Ighg1tm1(IRES-cre)Cgn/J (Cγ1^cre^) mice were originally purchased from The Jackson Laboratory and bred in house. B6.Cg-Tg(Fcer2a-cre)5Mbu (B6.CD23^cre^) mice were originally provided by Dr. Meinrad Busslinger (University of Vienna) which were crossed to STAT3^flox^ mice in house. Tg(MS4A1-cre/ERT2)1Mjsk (hCD20^ERT2-Cre^) mice were generously provided by Dr. Mark Shlomchik (University of Pittsburgh) and bred to STAT3^flox^ mice in house. All animal studies were conducted at Pennsylvania State University Hershey Medical Center in accordance with the guidelines approved by our Institutional Animal Care and Use Committee. Animals were housed in a specific pathogen-free barrier facility.

### Immunization and Influenza Viral Infection

For the studies of the peak GC response, GC maintenance, and long-lived plasma and memory B cell responses, we employed a prime-boost approach in which 8-10 wk old age and sex matched mice were immunized intraperitoneally (i.p.) with 200µg of 4-Hydroxy-3-nitrophenylacetyl-Keyhole Limpet Hemocyanin (NP-KLH, Biosearch Technologies) mixed with Complete Freund’s Adjuvant (CFA, Sigma-Aldrich) on d0 followed by 100µg NP-KLH in Incomplete Freund’s Adjuvant (IFA, Sigma-Aldrich) on d7. For the studies of early GC response or GC initiation, 8-10 wk old mice were immunized i.p. with 200µg of NP-KLH in CFA only. Spleens were harvested and processed into a single cell suspension for various analysis on indicated day post immunization. Influenza viral infection was performed by infecting mice intranallasy with 1000 fluorescent focus units of the H1N1 strain (A/Puerto Rico/8/34) in a 40µL volume as described (Fino et al., 2017). Mice were anesthetized using inhaled isoflurane before intranasal inoculation. Weight loss was tracked by daily measurements. Mediastinal lymph nodes (mLN) were harvested from infected animals on d14 post-viral infection and processed for various downstream analysis.

### Statistical Analysis

All results represent the mean of the data ± Standard Error of the Mean (SEM), unless indicated otherwise. For all experiments the normality of the dataset was first determined using D’agostino-Pearson normality test. Statistical significance between two groups was then determined using the Student’s *t*-Test or Mann-Whitney U when two groups were compared or an ANOVA when more than two groups were analyzed. The statistical test used is denoted in each figure legend. Statistical analyses were performed using Prism GraphPad software version 6 (GraphPad Software). P values are shown as *, p <0.05, **, p <0.01, ***, p <0.001, ****, p <0.0001 for significance.

## Acknowledgments

We would like to thank the Pennsylvania State University Hershey Medical Center flow cytometry core facility for help with flow cytometric experiments. We would also like to thank the Pennsylvania State University Hershey Medical Center Department of Comparative Medicine for help with animal housing and care. We thank the support staff within the central facility of the Department of Microbiology and Immunology at Pennsylvania State University College of Medicine. We would like to acknowledge the contribution of Vivian Gersuk and the Genomics Core at Benaroya Research Institute with their help with RNAseq. Further, we thank Dr. Daisuke Kitamura for kindly providing 40LB fibroblast cell line. Finally, we would like to thank Drs. Zissis Chroneos for the influenza virus and Aron Lukacher for the critical reading of the manuscript.

## Funding

National Institutes of Health grant R01 AI162971 (ZSMR)

National Institutes of Health grant R01AI091670 (ZSMR)

National Institutes of Health grant R01AI143778 (MRC)

Lupus Research Alliance Award 548931 (ZSMR)

Finkelstein Memorial Award (PPD)

National Cancer Institute 5T32CA060395-25 (AJF)

National Center for Advancing Translational Sciences UL1TR002003 (MM)

## Author contributions

Conceptualization: AJF, SBC, ZSMR

Methodology: AJF, SBC, ZSMR

Investigation: AJF, SBC, KNB, JLW, SLB, NC, NEW, MM

Formal Analysis: AJF, NEW, MM, AR, MM, PPD, ETLP, ZSMR

Funding acquisition: ZSMR

Resources: MRC

Writing-original draft-AJF, ZSMR

Writing-review & editing-AJF, ZSMR

## Competing interests

The authors have no financial conflicts of interests to disclose.

## Notes

### Competing Interest Statement

The authors have declared no competing interest.

