## Supplemental materials and Methods for "STAT3 signaling in B cells controls germinal center zone organization and recycling"

**Supplementary Materials and Methods**

**Flow Cytometry**

Spleens or mediastinal lymph nodes were harvested from indicated mice at the indicated time points and processed into a single cell suspension followed by RBC lysis using Tris-Ammonium Chloride. Cells were then stained with various antibodies described in the table below. To examine the endogenous levels of phosphorylated STAT3 in GC B cells, we used a previously characterized system where tissues are immediately processed into a single cell suspension in a fixative (Khalil et al., 2012). In brief, harvested tissues were immediately processed in 1.5% paraformaldehyde (Thermo Fisher Scientific) diluted in RPMI and incubated for 15 minutes at room temperature. Fixed cells were then washed once with PBS followed by RBC lysis prior to staining using antibodies in the table below. For intracellular staining, cells were fixed using BD Cytofix (BD Biosciences) and permeabilized using BD Phosflow Perm Buffer III according to manufacturer's instructions. All staining of unfixed cells was performed at 4□C to prevent internalization except for anti-CXCR5 which was performed at room temperature. Stained cells were analyzed on a BD LSR II or BD FACSymphony A3 (BD Biosciences) using FACSDiva Software. Sorting experiments were performed on a BD ARIA SORP high-performance cell sorter (BD Biosciences). Flow cytometry data was analyzed using FlowJo Software version 10.5.0 (BD).

| **Marker** | **Conjugate/Dye** | **Clone** | **Manufacturer** |
|---|---|---|---|
| B220 | BV605 | RA3-6B2 | Biolegend |
| T-and B-Cell Activation Antigen | FITC | GL-7 | BD Biosciences |

| | | | |
|---|---|---|---|
| T-and B-Cell Activation Antigen | PE | GL-7 | BD Biosciences |
| CD95 | PE-Cy7 | Jo2 | BD Biosciences |
| CXCR4 | BV711 | 2B11 | BD Biosciences |
| CD86 | PE-Cy5 | GL-1 | Biolegend |
| CD86 | PE | GL1 | BD Biosciences |
| CD138 | PE-Cy7 | 281-2 | Biolegend |
| CD138 | APC | 281-2 | Biolegend |
| CD138 | BV421 | 281-2 | Biolegend |
| TACI | PE | 8F10 | Biolegend |
| TACI | Alexa Fluor 647 | 8F10 | BD Biosciences |
| IgD | APC | 11-26c.2a | BD Biosciences |
| B220 | Pacific Blue | RA3-6B2 | BD Biosciences |
| CD19 | BV605 | 6D5 | Biolegend |
| CD4 | Alexa Fluor 700 | RM4-5 | Biolegend |
| CD4 | PE | GK1.5 | BD Biosciences |
| CD44 | APC | IM7 | Biolegend |
| CD62L | PE-Cy7 | MEL-14 | Biolegend |
| CXCR5 | Biotinylated | 2G8 | BD Biosciences |
| PD-1 | PE | 29F.1A12 | Biolegend |

| | | | |
|---|---|---|---|
| IgD | Alexa Fluor 700 | 11-26c.2a | Biolegend |
| Bcl-6 | V450 | K112-91 | BD Biosciences |
| STAT3 | APC | M59-50 | BD Biosciences |
| STAT3 Phospho-Serine727 | PE | 49/p-Stat3 | BD Biosciences |
| STAT3 Phospho-Tyrosine 705 | PE | 4/P-STAT3 | BD Biosciences |
| BrdU | APC | 3D4 | BD Biosciences |
| EdU | Alexa Fluor 488 | N/A | Thermo Fisher Scientific |
| Hoescht (33342) | N/A | N/A | Thermo Fisher Scientific |
| Ephrin-B1 | Biotinylated | Polyclonal | R&D Systems |
| CD38 | BUV395 | 90/CD38 | BD Biosciences |
| CD38 | FITC | 90 | Biolegend |
| CD69 | Alexa Fluor 700 | H1.2F3 | Biolegend |
| NP | PE | N/A | Biosearch Technologies |
| FOXO1 | PE | C29H4 | Cell Signaling Technologies |
| cMYC | APC | 9E10 | Novus Bio |
| Cyclin D3 | PE | DCS-22 | Biolegend |
| Phospho-S6 (Ser235/236) | Pacific Blue | D57.2.2E | Cell Signaling Technologies |

| | | | |
|---|---|---|---|
| Phospho-Akt (Ser473) | Alexa Fluor 647 | D9E | Cell Signaling Technologies |
| IgG1 | Biotinylated | Polyclonal | Invitrogen |
| IgG1 | PE | A85-1 | BD Biosciences |
| FDC-M1 | Unconjugated | FDC-M1 | BD Biosciences |
| Ki-67 | PE | 16A8 | Biolegend |
| Phospho-H2A.X (Ser139) | APC | 2F3 | Biolegend |
| CD154 | Unconjugated | MR-1 | Biolegend |
| Fixable Viability Dye | eFluor780 | N/A | Invitrogen |
| Annexin V | APC | N/A | Invitrogen |
| STAT3 | Unconjugated | D3Z2G | Cell Signaling Technologies |
| Streptavidin | PE-Cy5 | N/A | Biolegend |
| Streptavidin | PerCP-Cy5.5 | N/A | eBioscience |
| Streptavidin | v500 | N/A | BD Biosciences |
| Streptavidin | APC | N/A | Invitrogen |

***In vivo* Treatments**

To study memory B cell development, 1mg of Bromodeoxyuridine (BrdU, Sigma) was injected intraperitoneally every 12 hours on d11-13 post-NP-KLH immunization as described (Weisel et al., 2016). *In vivo* BrdU pulse for proliferation and BrdU/5-ethyl-2’-deoxyuridine (EdU, Sigma) dual pulse for cell cycle analysis were conducted as previously described (Gitlin et al., 2014). In short, for single BrdU injection, mice were pulsed with 2mg of BrdU (i.v.) and harvested 90 minutes later. For the BrdU/EdU dual pulse, mice were first injected with 2mg of BrdU (i.v.) then pulsed an hour later with 1mg of EdU (i.v.). Mice were harvested thirty minutes after the EdU pulse. For flow cytometric detection of EdU, surface staining of cells was performed followed by Click-iT™ EdU Alexa Fluor™ 488 Flow Cytometry Assay Kit according to manufacturer's instructions (Thermo Fisher Scientific). Detection of BrdU was performed using the APC BrdU Kit according to manufacturer's instructions (BD Biosciences). B:T blockade was performed by the administration of 200μg of Ultra-Leaf purified anti-CD154 (clone MR1) (Biolegend) in PBS (i.v.) 12 hours prior to harvest on d14 post immunization. Control animals received an equivalent volume of PBS. Tamoxifen administration to hCD20$^{ERT2\text{-}Cre}$ mice was performed as previously described on indicated days (Khalil et al., 2012; Luo et al., 2018). Mice received 100μL of 10mg/mL solution of tamoxifen (Sigma) dissolved in corn oil (Sigma) by oral gavage.

**ELISA**

High affinity (NP4) and total (NP29) NP-specific serum antibody titers were measured as previously described (Fike et al., 2021). In brief, Immulon 4 HBX Microiter Plates (Thermo Fisher Scientific) were coated with 10μg/mL of $NP_4$-BSA or $NP_{29}$-BSA and blocked with 5% BSA in PBS. Serum was then added followed by serial dilution. For the detection of influenza virus-specific antibodies, ELISA plates were coated with 175ng of influenza virus and blocked with 5% BSA in PBS. Serum was added to the plate beginning with a 1:25 dilution, followed by serial dilution. Abs were then detected with one of the following biotinylated Abs: goat anti-mouse IgM (Jackson Immunoresearch), goat anti-mouse IgG (Jackson Immunoresearch), goat anti-mouse IgG1 (Invitrogen), or goat anti-mouse IgE (Southern Biotech), followed by streptavidin-alkaline phosphatase (Vector Laboratories). Plates were developed using p-nitrophenyl phosphate (disodium salt) (Thermo Fisher Scientific) and read at λ405nm on a Synergy H1 (BioTek Instruments).

**ELISpot**

To detect NP-specific antibody-forming cells (AFCs), Hydrophobic High Protein Binding ELISpot plates (Millipore Sigma) were coated with 10μg/mL $NP_4$-BSA or $NP_{29}$-BSA and blocked with 5% BSA in PBS. 0.2 $x10^6$ splenocytes or bone marrow cells were serially diluted and cultured overnight at 37°C in RPMI with 10% FBS (Corning), 1X glutamax (Corning), and 1X Penicillin-Streptomycin (Corning). AFCs were detected with biotinylated goat anti-mouse IgG (Jackson Immunoresearch) or goat anti-mouse IgG1 (Invitrogen) Abs followed by streptavidin-alkaline phosphatase (Vector Laboratories). Plates were developed with Vector Blue Alkaline Phosphatase Substrate Kit III (Vector Laboratories). ELISpots were enumerated using an ELISpot plate imaging/analysis system ImmunoSpot S6 Universal (Cellular Technology Limited).

**Immunofluorescence Microscopy**

Splenic tissue was first embedded in Tissue-Tek O.C.T. compound (Sakura) and snap frozen over liquid nitrogen in 2-Methylbutane (Fisher Scientific). Five-μm sections were cut using a cryostat (Hacker Instruments and Industries), mounted on ColorFrost Plus Microscope Slides (Thermo Fisher Scientific) and fixed in cold acetone for 20 minutes. Sections were stained with the following Abs: GL-7-FITC (GL-7; BD Biosciences), CD4-PE (GK1.5; Biolegend), IgD-APC (11-26c2a; BD Biosciences), CD86- PE (GL-1; BD Biosciences), IgG1-Biotinylated (Invitrogen), or FDC-M1- Purified (FDC-M1; BD Biosciences). For active caspase 3/7 detection, we performed FAM-FLICA on unfixed splenic sections according to manufacturer instructions (ImmunoChemistry Technologies).

***In vitro* GC Culture**

*In vitro* GC (iGC) culture was performed as described (Nojima et al., 2011). B cells were isolated from indicated mice using a negative selection B cell isolation Kit (StemCell Technologies). B cells were overlaid on irradiated (120gy) 40LB fibroblasts in RPMI (Corning) containing 10% Hyclone Fetal Bovine Serum (GE Life Sciences), Penicillin Streptomycin (Corning), Glutamax (Gibco), Sodium Pyruvate (Gibco), 5μM 2-Mercaptoethanol, and HEPES Buffer (Corning) supplemented with 5ng/mL IL-4 (Peprotech) at 37C in 5% CO2. On day 4 of the culture GC B cells were purified from the fibroblasts by negative selection using biotinylated H-2Kd (SF1-1.1; Biolegend) and CD138 (281-2; BD Biosciences) Abs and stimulated with various concentrations of IL-21 (Peprotech). Cells were harvested 12 hours post-stimulation and processed for RT-PCR as described below.

**VH186.2 BCR sequencing and alignment**

Mutation analysis of VH186.2 was performed as described (Rydyznski et al., 2018). NP-specific GC B cells from $CD23^{cre}$ and $STAT3^{fl/fl}CD23^{cre}$ mice were FACS sorted and frozen in Trizol (Life Technologies) on d14 post-immunization. Total RNA was isolated using the Direct-zol RNA Miniprep kit (Zymo Research) and was converted to cDNA using the High-Capacity cDNA Reverse Transcription Kit (Thermo Fisher Scientific). Two rounds of nested PCR were performed to specifically amplify the VH186.2 NP-specific heavy chain using GoTaq G2 Hot Start polymerase (Promega) and the following primers: Round 1 Fwd: CTCTTCTTGGCAGCAACAGC Rev: GCTGCTCAGAGTGTAGAGGTC, Round 2 Fwd: GTGTCCACTCCCAGGTCCAAC, Rev: GTTCCAGGTCACTGTCACTG. PCR amplified sequences were then agarose gel purified using a 1% gel and DNA was isolated using the QIAquick Gel Extraction Kit (QIAGEN). Purified DNA was ligated into the pGEM-T easy vector (Promega) and used to transform XL-10 Gold ultracompetent *E. coli* (Agilent) and plated on Xgal-IPTG-Amp eight agar plates (1 per mouse) (KD Medical). Plates were incubated overnight at 37□C. The following day, six white colonies per plate (24 colonies per genotype) were selected at random and sequenced by Genewiz. Sequences were then aligned to the germline using NCBI IgBlast browser as described (Rydyznski et al., 2018). Total number of mutations relative to the VH186.2 germline were recorded, along with their location, type of mutation (silent vs replacement), and presence or absence of the tryptophan to leucine (W33L) mutation.

**BCR repertoire analysis**

$CD23^{cre}$ and $STAT3^{fl/fl}CD23^{cre}$ mice were immunized with NP-KLH as described in Materials and Methods. Mice were sacrificed 21 days after initial immunization. $NP^{+}$ and $NP^{-}$ GC B cells were FACS purified from splenocytes pooled from 4 mice of each genotype. RNA isolation, amplification of BCR heavy chains, and sequencing was performed by iRepertoire Inc. FASTQ files from iRepertoire were processed for downstream analysis first using pRESTO (Vander Heiden et al., 2014) to assemble paired reads, remove low quality and short reads, and mask low quality bases as described in(Kuri-Cervantes et al., 2020). Next, reads were aligned and annotated with gene calls using IgBLAST (Ye et al., 2013) and imported into ImmuneDB(Rosenfeld et al., 2018). Sequences with the same V-gene, J-gene, and CDR3 length were hierarchically clustered and those sharing at least 85% amino-acid homology in the CDR3 were grouped into clones. All further analyses only included productive clones.

**RT-PCR**

RNA from in vitro generated GC B cells was isolated using the Qiagen RNeasy kit according to manufacturer's instructions (Qiagen) after removing any contaminating fibroblasts as described above. RNA quality and quantity was assessed via nanodrop (Thermo Scientific) and cDNA was generated using the High-Capacity cDNA Reverse Transcription Kit (Thermo Fisher Scientific) according to manufacturer's instructions. *Foxo1* transcript levels were measured using the following primer sequences and SYBR Green Master Mix (Thermo Fisher Scientific): Forward: 5'-CCCAACCAAAGCTTCCCACA-3' and Reverse: 5'-AAATGTAGCCTGCTCACTAACTCT-3'.

Transcript levels were normalized against a house keeping gene (β-actin), and fold change was calculated via the $2^{(\Delta\Delta Ct)}$ method.

**RNA sequencing and Pathway Analysis**

Splenic $NP^+$ GC B cells (Live $B220^+GL\text{-}7^+CD95^+$) were sorted from $CD23^{cre}$ and $STAT3^{fl/fl}CD23^{cre}$ mice on 14d post-NP-KLH immunization for RNA sequencing. Total RNA was prepared from sorted cells using the Qiagen RNeasy kit according to manufacturer's instructions. Total RNA (0.5ng/mL) was added to lysis buffer from the SMART-seq V4 Ultra Low Input RNA kit for Sequencing (Takara), and reverse transcription was performed followed by PCR amplification to generate full-length amplified cDNA. Sequencing libraries were constructed using the NexteraXT DNA sample preparation kit (Illumina) to generate Illumina-compatible barcoded libraries. Libraries were pooled and quantified using a Qubit® Fluorometer (Life Technologies). Dual-index single-read sequencing of pooled libraries was carried out on a HiSeq2500 sequencer (Illumina) with 58-base reads using HiSeq v4 Cluster and SBS kits (Illumina) with a target depth of 5 million reads per sample. Base calls were processed to FASTQs on BaseSpace (Illumina), and a base call quality-trimming step was applied to remove low-confidence base calls from the ends of reads. Data were aligned to the GRCm38 reference genome using STAR v.2.3.2a and gene counts were generated using htseq-count. QC and metrics analysis was performed using the Picard family of tools (V1.134). Gene set enrichment plots were generated from total gene counts using GSEA (version 4.0.3), and the Principal Component Analysis was performed using R (Version 1.1.456). Differentially expressed genes between $CD23^{cre}$ or $STAT3^{fl/fl}CD23^{cre}$ GC samples were calculated with Limma (Version 3.42.2) using a fold-change cutoff of 1.5 and an adjusted p-value cutoff of 0.05. Volcano plots were generated using the ggplot2 package in R. Heatmap plots were generated from differentially expressed gene lists using Java Treeview (Version 1.1.6r4). Finally, Gene set enrichment plots were generated with the Gene Set Enrichment Analysis (GSEA) tool (Version 4.0.3) (Mootha et al., 2003; Subramanian et al., 2005).

**ChIP-seq**

For the analysis of STAT3 binding sites, C57BL/6 mice were immunized as described above. Spleens from 10 mice were harvested, processed into single cell suspensions, and pooled on 14d post immunization. ChIP-seq samples were generated from two experiments using a total of 20 mice. Following RBC lysis using Tris-Ammonium Chloride, B cells were first enriched by negative selection (Stem Cell Technologies) and then stained for GC B cell markers viability- , anti-B220 BV605, anti-GL7 FITC and anti-CD95 PE-Cy7. GC B Cells were sorted using a FACS Aria SORP high-performance cell sorter (BD Biosciences) and immediately cross-linked in 37% formaldehyde (Sigma) diluted in PBS (Corning) to a final concentration of 1.5% formaldehyde. Cells were fixed for 12 minutes at 37□C and then reaction was stopped by adding Glycine (Cell Signaling Technologies) and a 10 minute incubation at room temperature. Cells were then washed with PBS and the pellet was snap frozen and stored at -80□C until processing.

ChIP-seq was performed using the ChIP Assay Kit following the manufacturer's protocol (Millipore). DNA was sheared to ~200 bp with a Bioruptor sonication water bath for four 15 min cycles consisting of 45s on and 30s off. ChIP was performed with anti-STAT3 (D3Z2G, Cell Signaling Technologies). Resulting DNA was purified with the MinElute PCR Purification Kit (Qiagen). DNA was incubated with a tagmentation enzyme (Illumina) at 37 °C for 30 min and DNA was purified with the MinElute PCR Purification Kit (Qiagen). DNA libraries were made with the Nextera Indexing kit (Illumina), and DNA was purified with the QiaQuick PCR Purification Kit (Qiagen). Libraries were sequenced on the Illumina NovaSeq6000.

Raw sequence reads were aligned to reference genome mm9 using BWA MEM (Li, 2013). PCR duplicates were removed using Picard MarkDuplicates (https://github.com/broadinstitute/picard/releases/tag/2.11.0). Peaks were called against input using Macs2 (Zhang et al., 2008). Normalized bedgraph tracks were generated using the SPMR flag and converted to bigWig using the UCSC tool bedGraphToBigWig. Peaks with a score >50 were retained. Hypergeometric optimization of motif enrichment (HOMER) was used to perform known transcription factor binding motif analyses using the FindMotifsGenome function (Heinz et al., 2010).

STAT3 peaks were converted to mm10 coordinates using UCSC liftOver. From the $STAT3^{-/-}$ RNA-seq described above, differentially expressed genes with STAT3 ChIP peaks within 50 kb upstream or downstream of the transcription start site were identified. RNA expression of these genes across GC zones was previously published (Kennedy et al., 2020). Genes with <1 copy per million in LZ GC B cells were excluded. RNA expression was plotted using ComplexHeatmap in R. ChIP peaks were visualized with the Integrated Genome Browser.

**Fig. S1. The organization of GC DZ and LZ at an early stage of GC formation requires B cell intrinsic STAT3 signaling**

**Fig. S2. T cell intrinsic STAT3 deficiency impairs Tfh development, but not the DZ and LZ organization**

**Fig. S3. Somatic hypermutation, and BCR clonal diversity**

**Fig. S4. Proliferation and cell death are not affected in the absence of STAT3 in B cells**

**Fig. S5. Genes and pathways regulated by STAT3 in GC B cells**

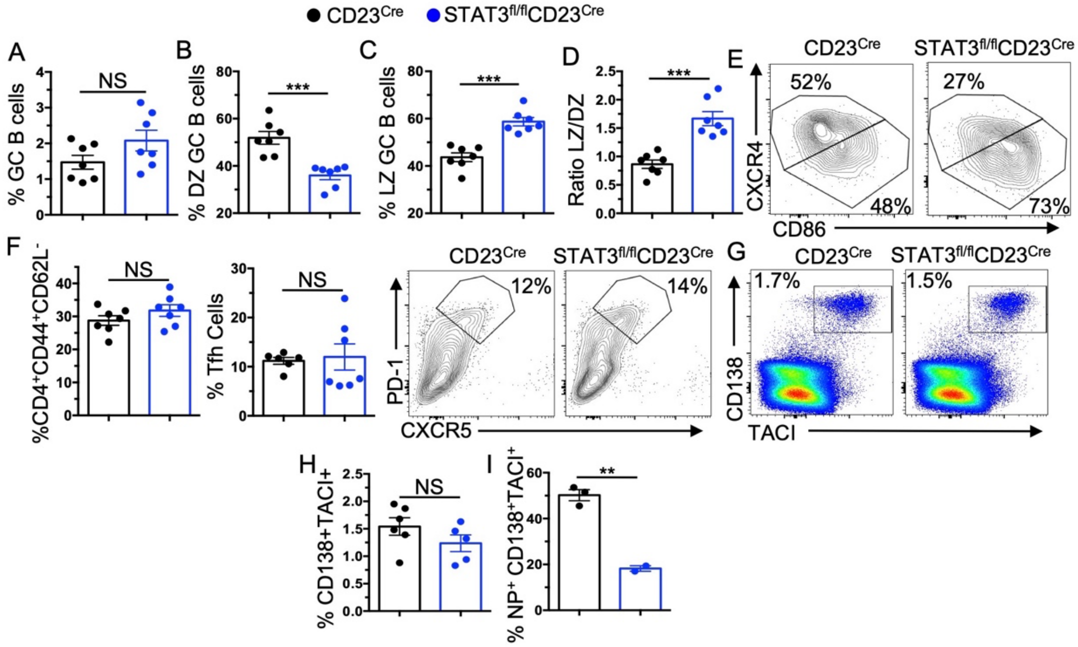


**Figure S1 - The organization of GC DZ and LZ at an early stage of GC formation requires B cell-intrinsic STAT3 signaling.** STAT3$^{fl/fl}$CD23$^{Cre}$ and CD23$^{Cre}$ control mice were immunized with NP-KLH/CFA and the splenic response was analyzed on day 7 post-immunization. (**A-E**) Flow cytometric quantification of GC, DZ and LZ B cell frequency, and representative flow plots of LZ/DZ organization. (**F**) Frequency of effector and Tfh cells and representative flow plots of Tfh populations. (**G-H**) Representative flow plots of plasma cell populations (IgD$^{-}$CD138$^{+}$TACI$^{+}$) and the frequency of total and NP-specific plasma cells. Each symbol represents an individual mouse and data are presented as means ± SEM (n=5-6 mice per group). Accumulative data represent three experiments. P values were calculated via an unpaired t-test or Mann-Whitney test (NS, not significant, **, p <0.01, ***, p <0.001).

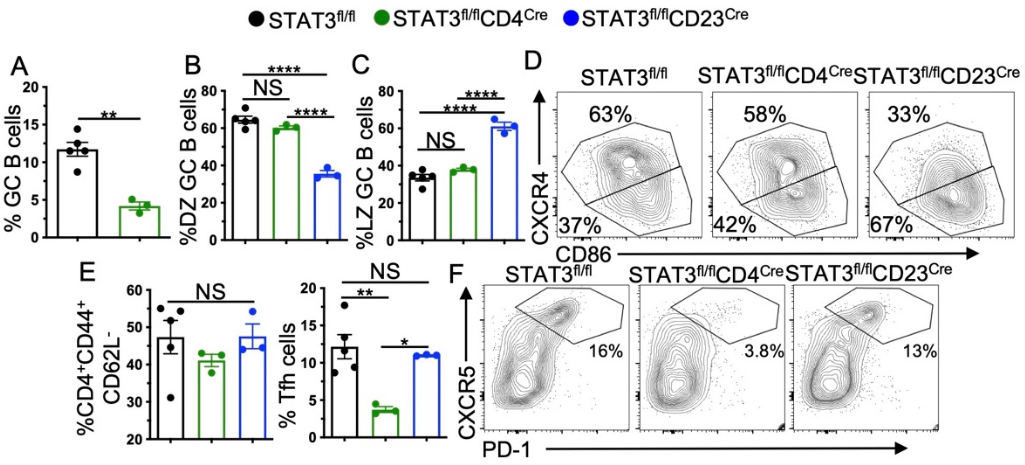


**Figure S2- T cell intrinsic STAT3 deficiency impairs Tfh development, but not the DZ and LZ organization**. **(A)** Percentage of splenic GC B cells in STAT3$^{fl/fl}$CD4$^{Cre}$ mice on 14d post-NP-KLH immunization. **(B-C)** Frequency of DZ and LZ GC B cells in indicated mice. **(D)** Representative flow plots of GC zone organization. **(E-F)** Frequency of effector CD4$^{+}$ T cells and Tfh in STAT3$^{fl/fl}$CD4$^{Cre}$, STAT3$^{fl/fl}$CD23Cre, and STAT3$^{fl/fl}$ control mice. **(F)** Representative flow plots of Tfh population pre-gated on CD4$^{+}$CD44$^{+}$CD62L$^{-}$. Each symbol represents an individual mouse and data are presented as means ± SEM (n=3-5 mice per group). Accumulative data represent two experiments. P values were calculated using a Mann-Whitney test or a One-Way ANOVA followed by Tukey's multiple comparisons test (NS, not significant, *, p <0.05 **, p <0.01, ****, p <0.0001).

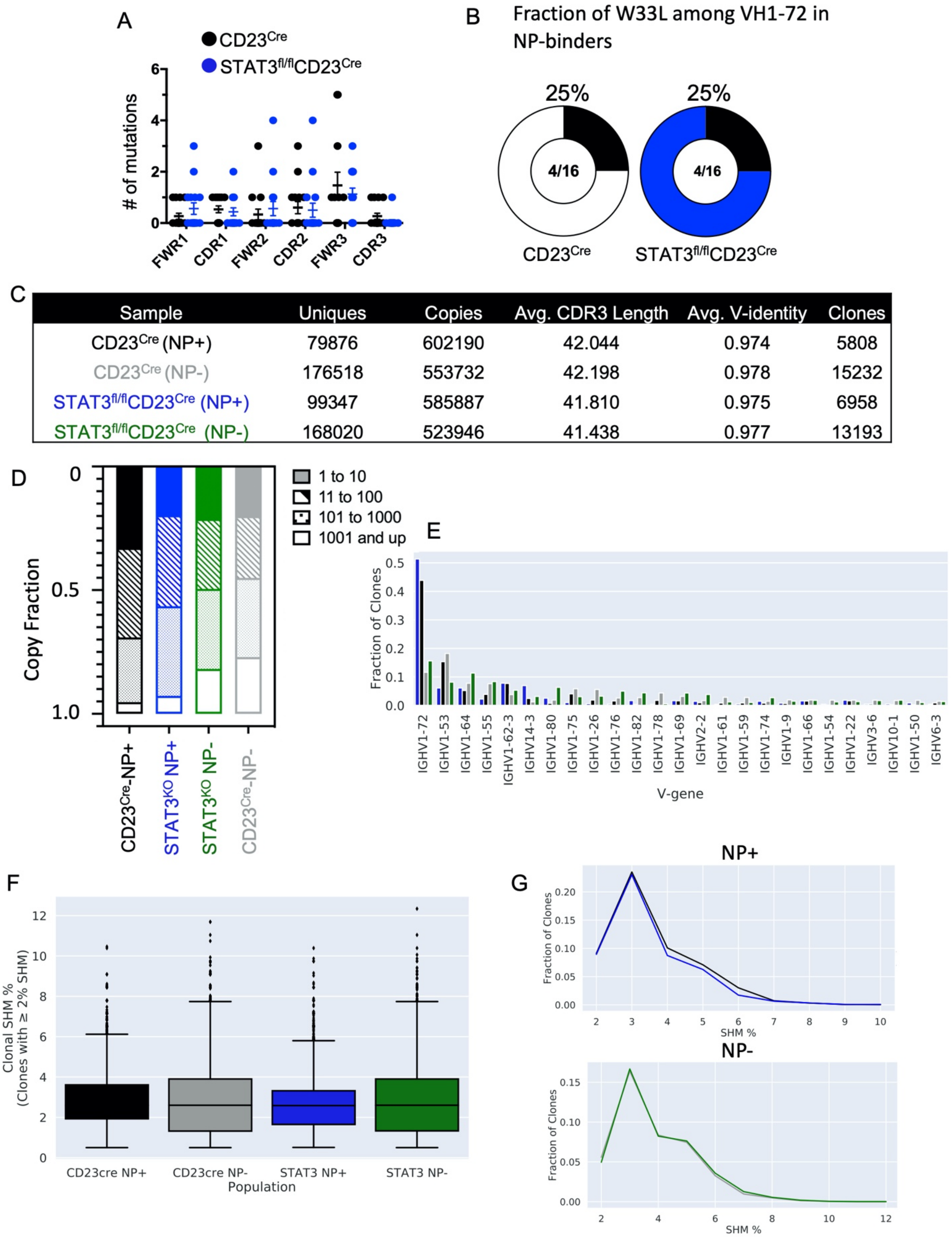


| Sample | Uniques | Copies | Avg. CDR3 Length | Avg. V-identity | Clones |
|---|---|---|---|---|---|
| CD23Cre (NP+) | 79876 | 602190 | 42.044 | 0.974 | 5808 |
| CD23Cre (NP-) | 176518 | 553732 | 42.198 | 0.978 | 15232 |
| STAT3fl/flCD23Cre (NP+) | 99347 | 585887 | 41.810 | 0.975 | 6958 |
| STAT3fl/flCD23Cre (NP-) | 168020 | 523946 | 41.438 | 0.977 | 13193 |

**Figure S3 - B cell intrinsic STAT3 deficiency exhibits minimal IgH repertoire differences compared to STAT3 sufficient B cells.** (**A-B**) Analysis of mutations within VH186.2 from NP$^{+}$

GC B cells sorted from CD23$^{cre}$ and STAT3$^{fl/fl}$CD23$^{cre}$ mice d14 post NP-KLH immunization by nested PCR and sanger sequencing. (**A**) Location of mutations in VH186.2 (FWR: Framework Region, CDR: Complementarity-determining region). (**B**) Of the 24 VH186.2 sequences mutations were identified in 16. Of the 16 sequences, 4 per group contained the W33L mutation (25%). Each symbol represents an individual sequenced bacterial colony and data are presented as means ± SEM (n=4 mice per group, 6 colonies/mouse, 24 colonies per group total) from two independent experiments. Only sequences which completely aligned with VH186.2 were considered. No data is statistically different. (**C-G**) Analysis of productive IgH rearrangement NGS data. (**C**) Metadata of sequencing libraries for STAT3 sufficient and deficient mice sorted on NP binding. Uniques indicates the total unique reads, Copies is the number of total reads, Avg. CDR3 Length is the mean CDR3 length in nucleotides, and Clones is the inferred number of clonotypes as defined in Methods. (**D**) Clonal range plots for each cell population. The fraction of reads in the top 10 copy-number clones is shown in solid colors, 11-100 ranked clones in hatch marks, 101-1000 in dots, and all smaller clones in white. (**E**) Fraction of VH-gene usage for the top-utilized 25 VH-genes in each of the four genotypes. (**F**) Boxplots of SHM for mutated clones (those with ≥ 2% mutation). (**G**) SHM distributions for each of the four populations. The top panel is for the NP$^{+}$ populations and the bottom panel for the NP$^{-}$ populations. P values were calculated using a Mann-Whitney U test (**, p<0.01, ****, p <0.0001).

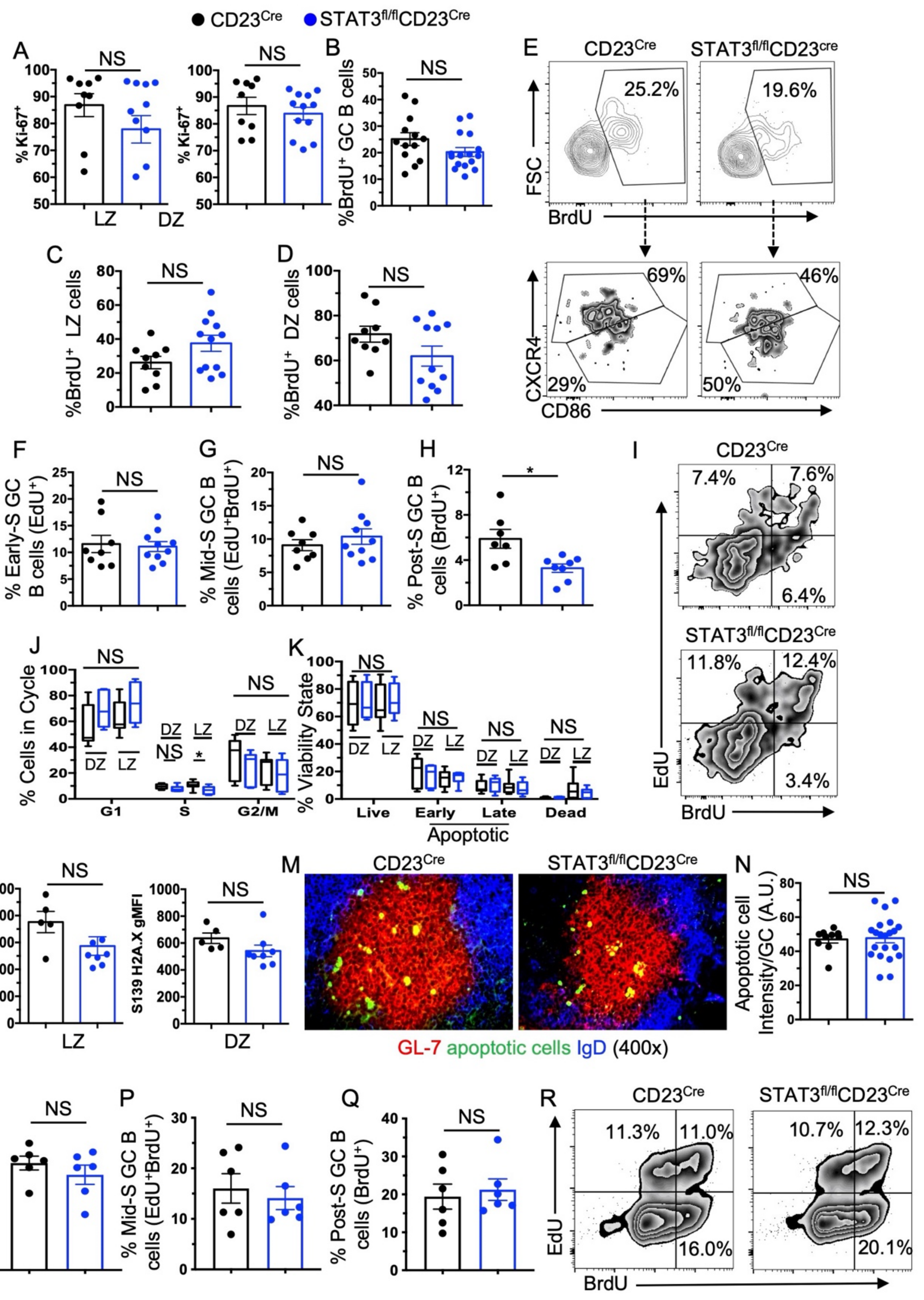


**Figure S4- Proliferation and cell death are not affected in the absence of STAT3 in B cells**. Analysis of proliferation, cell cycle, DNA damage, and apoptosis in CD23$^{Cre}$ and STAT3$^{fl/fl}$CD23$^{Cre}$ mice 14d post-NP-KLH immunization. **(A)** Frequency of Ki-67$^{+}$ LZ and DZ GC

B cells. **(B-D)** Analysis of BrdU$^+$ total GC, LZ, and DZ GC B cells 90 minutes after IV BrdU pulse. **(E)** Representative flow plots of BrdU$^+$ GC B cells and the respective LZ/DZ profile. **(F-I)** Analysis of S-phase progression in mice pulsed with BrdU IV followed by EdU IV 60 mins later and harvested 30 minutes post-EdU injection. Frequency of **(F)** early S-phase (EdU$^+$BrdU$^-$), **(G)** mid-S phase (EdU$^+$BrdU$^+$), and **(H)** post-S phase (EdU$^-$BrdU$^+$) GC B cells and **(I)** representative flow plots. **(J)** Cell cycle state quantified by DNA content in LZ and DZ GC B cells. **(K)** Quantification of live (Viability Dye$^-$Annexin V$^-$), early apoptotic (Viability Dye$^-$Annexin V$^+$), late apoptotic (Viability Dye$^+$ Annexin V$^+$), and dead cells (Viability Dye$^+$ Annexin V$^-$). **(L)** DNA damage assessed by gMFI of phosphorylated serine 139 of Histone H2A.X in LZ and DZ GC B cells. **(M)** Representative immunofluorescence microscopy image of GL-7$^+$ (Red) GCs and active Caspase 3/7$^+$ cells and **(N)** quantification of Caspase 3/7$^+$ cells. Frequency of **(O)** early S-phase, **(P)** mid-S phase, and **(Q)** post-S phase GC B cells and **(R)** representative flow plots on day 7 post-NP-KLH immunization. P values were calculated using an unpaired t-test or Mann-Whitney test (NS, not significant $p > 0.05$, *, $p < 0.05$).

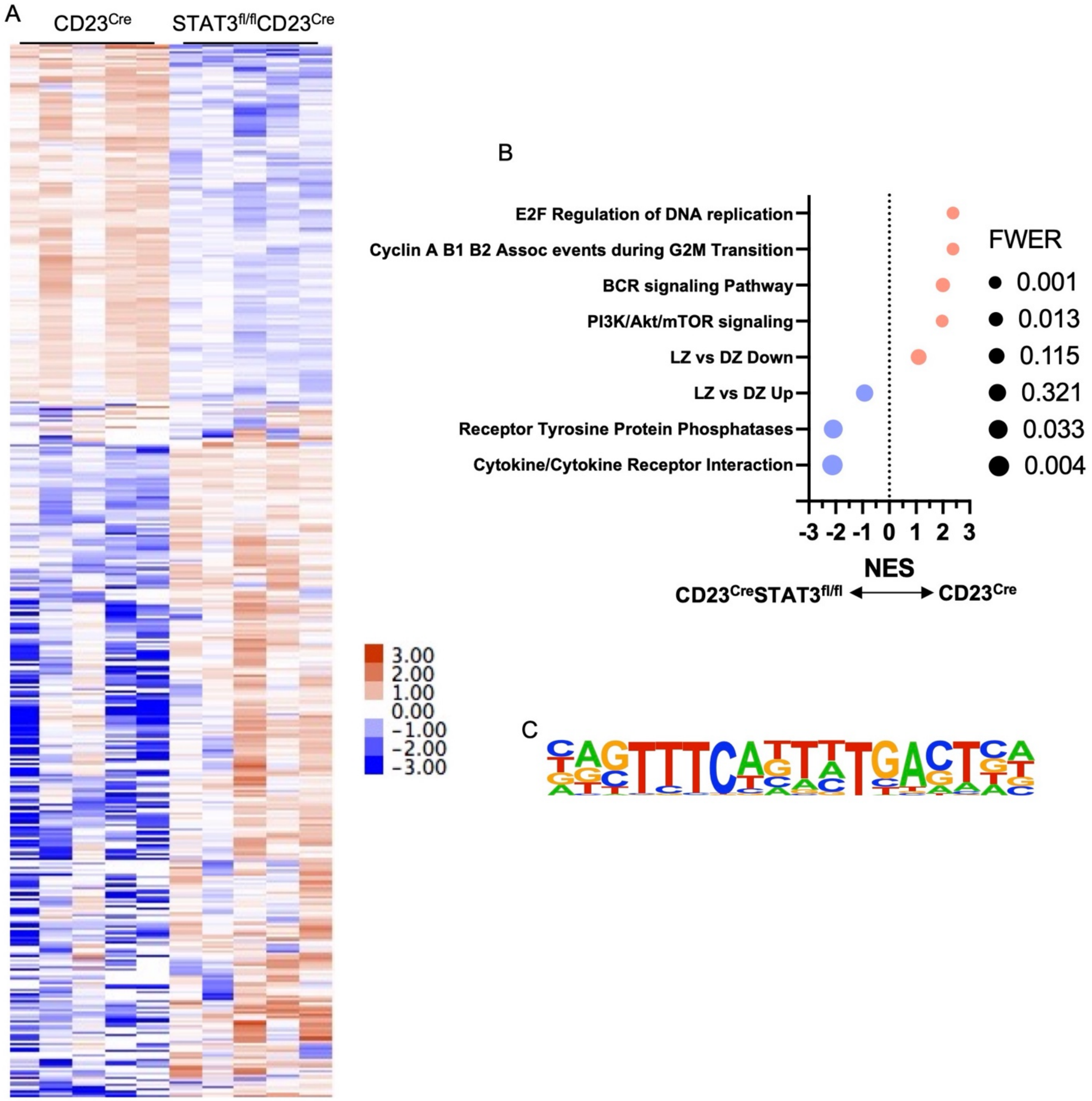


**Figure S5- Genes and pathways regulated by STAT3 in GC B cells**. **(A)** Heatmap of global transcriptional differences between CD23$^{Cre}$ and STAT3$^{fl/fl}$CD23$^{Cre}$ GC B cells on 14d post-NP-KLH immunization. **(B)** Selected GSEA pathways plotted by normalized enrichment score (NES) and their relative p-value adjusted by family-wise error rate (FWER). (**C**) Top motif of STAT3 ChIPseq peaks identified using HOMER ($p=10^{-744}$).
